# Humanized *in vivo* bone marrow models orchestrate multi-lineage human hematopoietic cell development

**DOI:** 10.1101/2024.04.08.588553

**Authors:** Laurent Renou, Wenjie Sun, Chloe Friedrich, Klaudia Galant, Cecile Conrad, Evelia Plantier, Katharina Schallmoser, Linda Krisch, Vilma Barroca, Saryami Devanand, Nathalie Déchamp, Andreas Reinisch, Jelena Martinovic, Alessandra Magnani, Lionel Faivre, Julien Calvo, Leila Perie, Olivier Kosmider, Françoise Pflumio

## Abstract

Hematopoiesis develops in the bone marrow (BM) where multiple interactions regulate differentiation and preservation of hematopoietic stem/progenitor cells (HSPCs). Although murine BM has been extensively analyzed, the human BM microenvironment remains less understood. Immune-deficient murine models have enabled the analysis of molecular and cellular regulation of human HSPCs, which remains limited as human hematopoietic cells develop in xenogenic microenvironments. In this study, we thoroughly characterized a humanized (h) *in vivo* BM model, based on mesenchymal stromal cell (MSC) differentiation (called hOssicles (hOss)), and hematopoietic cell compartments generated 3 months post-transplant of CD34^+^ cells using single-cell RNA sequencing and cellular barcoding. Serial isolation of MSCs and HSPCs from hOss and transplant experiments revealed the dynamic nature of these hBM niches. hOss altered human hematopoietic development by modulating myeloid/lymphoid cell production and HSPC levels. Clonal tracking highlighted hematopoietic cell cross-talks between the murine BM and hOss, indicating the multipotent or more restricted lineage origin of human hematopoiesis shared in the BM sites.

## Introduction

Post-natal hematopoiesis takes place in the bone marrow (BM), where numerous cell interactions promote the production of mature blood cells. Analysis of the BM microenvironment in murine models has enabled the detailed characterization of spatially distinct niches of the various types of hematopoietic stem and progenitor cells (HSPCs), including dormant/quiescent and activated hematopoietic stem cells (HSCs) (Ding and Morrison, 2013; Cordeiro Gomes et al., 2016; Pinho et al., 2018). Furthermore, single-cell RNA sequence analysis of BM niches has recently revealed the heterogeneous nature of non-hematopoietic cells that support hematopoietic cell production (Tikhonova et al., 2019; Baryawno et al., 2019; Baccin et al., 2020).

Studying the human BM microenvironment is more complicated than murine BM as it involves analysis of BM biopsy sections, which enables only general characterization but not genetic modification of the different cell components, thus greatly limiting the investigation of cell function. Humanized (h)BM models, which are generated *ex vivo* and *in vivo* by transplanting human BM cell components into immune-deficient mice, serve as valuable tools in biomedical research (Abarrategi et al., 2018; Dupard et al., 2020). These models are designed to replicate the complex cellular interactions that occur in the human BM microenvironment. By recapitulating the physiological conditions, these models provide researchers with a powerful platform to study hematopoiesis, immune cell development, and disease progression. hBM models incorporate components critical for hematopoiesis, such as stromal cells, and extracellular matrix components (Chen et al., 2012; Antonelli et al., 2016; Groen et al., 2012; Abarrategi et al., 2017; Fritsch et al., 2018; Reinisch et al., 2016). Inclusion of human endothelial cells that contribute to blood vessel development and are key supportive cells of hematopoiesis in the BM (Ramasamy et al., 2016; Itkin et al., 2016) has been successfully described *in vivo* (Chen et al., 2012; Passaro et al., 2017; Abarrategi et al., 2017). Although hBM models are chimeric with murine and human cell components, these systems enable the study of normal and pathological conditions, including hematological disorders, leukemia and immunodeficiency. Furthermore, these models which are based on mesenchymal stromal/stem cells (MSCs) and hematopoietic cell differentiation, offer the opportunity to study the effects of various factors, such as infection, therapeutic interventions and response, and genetic mutations that can be incorporated into HSCs or progenitors using CRISPR-Cas9 editing. By providing a controlled and reproducible system, these models enable researchers to gain extensive insight into the mechanisms underlying diseases and test potential therapies with greater precision. Moreover, the application of hBM models has proven valuable in cancer research, enabling the analysis of tumor progression, including infiltration and metastasis, in the context of a physiologically relevant microenvironment (Martine et al., 2017; Thibaudeau et al., 2014; Grigoryan et al., 2022). By providing a platform to evaluate the efficacy and toxicity of anti-cancer drugs, hBM models also facilitate the development of personalized treatment strategies.

It has been hypothesized that species-specific HSC microenvironment interactions are likely vital for human hematopoietic development (Holzapfel et al., 2015). However, the impact of humanized niches on human hematopoietic cell recovery generated following CD34^+^ HSPC transplant has not been assessed (Abarrategi et al., 2018). In this study, we aimed to thoroughly characterize a robust and reproducible human ossicles (hOss) hBM model adapted from previous studies (Reinisch et al., 2017a). We generated hOss by using human fetal (F) and post-natal (P-N) BM-derived MSCs transplanted subcutaneously into immune-deficient mice. We found that F/hOss and P-N/hOss reproducibly supported human hematopoietic cell development following peripheral injection of human CD34^+^ umbilical cord blood cells. Using flow cytometry and single-cell RNA sequencing, characterization of mature and immature human hematopoietic cells developing in hOss revealed a myeloid cell bias at the expense of B lymphoid cells, contrary to that observed in the BM of gold standard NOD scid gamma (NSG) mouse models (Pflumio et al., 1996; Henry et al., 2020), thus more accurately recapitulating human BM cells. Importantly, recovered hOss contained hMSCs capable of secondary hOss formation and human hematopoietic support, indicating that hOss are dynamic BM structures that maintain functional MSCs. Furthermore, hOss better supported immature human cells capable of secondary hematopoietic reconstitution after transplant. Finally, using a cellular barcoding tracking strategy, we found that many human hematopoietic cell clones generated in mice harboring hOss were myeloid biased. Most cell clones were multipotent and produced human hematopoietic cells in both hOss and murine BM (mBM), thus indicating cross-talk between BM sites. The dynamic hOss structure created a favorable environment for human myelopoiesis and multipotent HSPCs.

## Results

### Development of hOss originating from human bone marrow-derived fetal and post-natal hMSCs

hOss formation was carried out according to a previously developed protocol (Reinisch et al., 2017a). Briefly, we first isolated primary hMSCs from F/BM and P-N/BM after cell adhesion on plastic plates. hMSCs were amplified until confluent and split; a portion of the cells were stored frozen at passage 0, while the remaining cells were further amplified for <4 passages for downstream analyses in accordance with the protocol by Passaro et al. (Passaro et al., 2017). Kinetic analysis of cell growth indicated enhanced expansion of the majority F/hMSCs (4 of 5) compared with P-N/hMSCs (**Figure 1A**). Flow cytometry analysis of conventional hMSC markers, such as CD90, CD73, CD105 and CD44, and pan-hematopoietic CD45 and myeloid CD14/CD15 markers and endothelial CD31 confirmed the non-hematopoietic mesenchymal origin of the isolated adherent cells of F/BM and P-N/BM origin (**Figure 1B and SupFig1A**). No major phenotypic differences were observed between F/hMSCs and P-N/hMSCs. These hMSCs were capable of *ex-vivo* multilineage differentiation into adipogenic, osteogenic and chondrogenic cells upon exposure to the appropriate cell culture media (**SupFig1B**). To test the ability to generate hOss, a mixture of hMSCs, cold-matrigel, human platelet lysate and hBMP-7 was subcutaneously injected into immune-deficient NSG mice that were treated 5 days per week for 4 weeks with parathyroid hormone (to enhance hOss formation and growth) (**Figure 1C**) (Reinisch et al., 2017b; Martine et al., 2017). OsteoSense labeling and tomography analysis, which typically indicates regions with high bone turnover, showed bone remodeling in hOss derived from F/hMSCs and P-N/hMSCs (**SupFig1C**) (Lambers et al., 2013; Martine et al., 2017). At 8 weeks post-hMSC implantation, hOss were recovered from grafted mice and analyzed. The F/hMSC-derived hOss were larger and 5 times heavier (median: 579 mg) than P-N/hMSC-derived hOss (median: 94 mg) (**Figure 1D-E**). Murine hematopoietic Ter119^+^ and CD45^+^ cells were detected in F/hOss and P-N/hOss, albeit at slightly lower levels compared with mBM, thus showing that murine blood cells perfused the implanted hBM (**Figure 1F**). Histological analysis of hOss sections indicated the presence of active bone structures, including adipocytes and hematopoietic lodges, resembling BM (**Figure 1G**). Immuno-labeling with anti-mouse endomucin or anti-mouse Meca32 antibodies also revealed vascular networks (**SupFig1D-E**), thereby confirming that murine blood vessels vascularized hOss. Overall, these results show efficient *in vivo* development of vascularized functional BM structures derived from human F/hMSCs and P-N/hMSCs.

**Figure 1.**
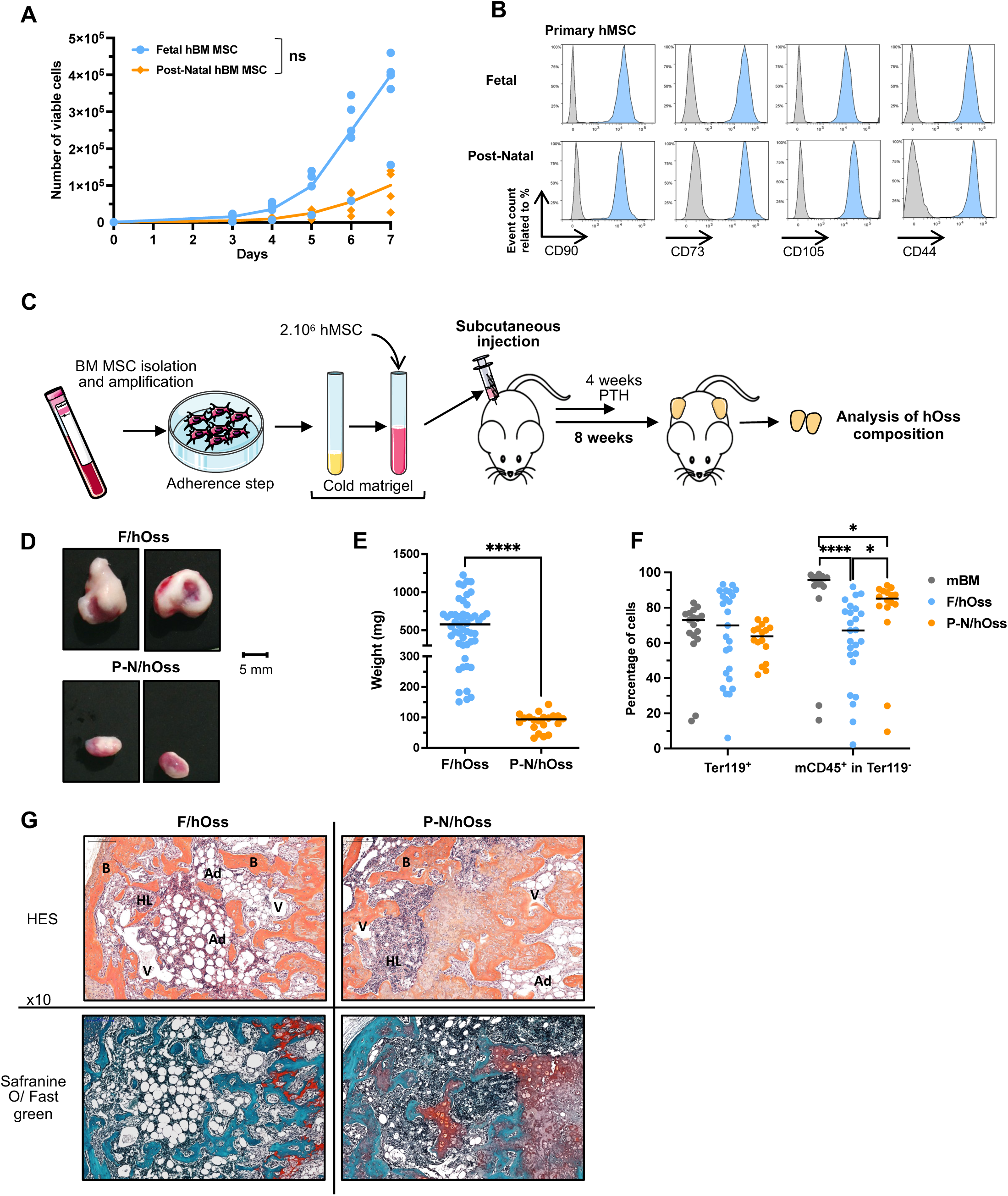
Human MSCs from fetal and post-natal bone marrow generates hOss. A. Growth curve of hMSCs isolated from fetal bone marrow (BM) (5 hMSCs, age <12 post-conception weeks (PCWs), passage 2) and post-natal BM (4 hMSCs, ages: 5, 9, 24 and 51 years, passage 2). Each culture was carried out in quadruplicate. The mean number of cells for each time point for every tested MSC is indicated; the median growth of fetal (in blue) and post-natal (in orange) MSCs are indicated. ns, not significant; multiple Mann-Whitney test. B. Phenotype analysis of adherent fetal (F28 sample, 11.6 PCWs, passage 2) and post-natal (ALLO3 sample, 5 years old, passage 2) human BM (hBM) cells isolated from fetal and post-natal BM. Cells were gated on hCD45 CD14 CD15 CD31 cells (>90%). Representative of >5 tested hMSC samples. The results were obtained from the gating strategy shown in SupFig1A. C. Experimental protocol of hOss generation using immune-deficient NSG mice. hMSCs were isolated from human BM samples obtained before (fetal, F) and after (post-natal, P-N) birth and amplified in plastic wells (passages ≤2), before mixing 2.10^6^ cells with 300µl cold-matrigel containing human platelet lysate (50µl) and hBMP7 (5µg/5µl) and subcutaneous transplantation in mice. Eight weeks later, hOss were harvested and analyzed. D. Representative pictures of hOss generated from fetal (F/hOss, F28, 11.6 PCWs, passage 1) and post-natal (P-N/hOss, ALLO2, 9 years old, passage 1) hMSCs. hOss were recovered 8 weeks after implantation in mice. E. Weight comparison of F/hOss and P-N/hOss. The results are shown from F/hOss (9 independent experiments, 11 hMSC samples; 50 hOss) and P-N/hOss (4 independent experiments, 5 hMSC samples; 20 hOss). Each dot represents a single hOss, and the line indicates the median value. ****, p<0.0001, Mann-Whitney test. F. Flow cytometry analysis of murine cells recovered from hOss. Percentage of cells in SSC/FSC gated viable cells are shown. Murine red blood cells are Ter119^+^, and murine hematopoietic cells are Ter119-/mCD45^+^. Results were obtained from F/hOss (4 independent experiments, 8 hMSC samples; 25 hOss) and P-N/hOss (3 independent experiments, 4 hMSC samples; 16 hOss). Lines indicate the median value of each condition. *, p<0.05; ****, p<0.0001; Kruskal-Wallis test without correction. G. Representative sections of F/hOss (left) and P-N/hOss (right) with histological staining using HES (upper panels) and Safranin O / Fast Green (lower panels). HL, hematopoietic lodge; V, vessel; B, bone; Ad, adipocytes. Amplification ×10.

### Human hematopoiesis in hOss after transplantation of human HSPCs

We next aimed to analyze the supportive ability of F/hOss and P-N/hOss for human hematopoiesis following intravenous transplantation of human CD34^+^ cells (10^5^ cells/mouse) in sub-lethally irradiated mice harboring hOss (**Figure 2A**). Human hematopoietic development was analyzed 12 weeks later, which is representative of long-term hematopoietic reconstitution in transplant models (Dykstra et al., 2007). We found that the size and weight differences persisted between F/hOss (median: 390 mg, at 12 weeks) and P-N/hOss (median: 78 mg, at 12 weeks) **(SupFig2A**) as observed in non-transplanted hOss (**Figure 1E**). Immunohistochemistry labeling indicated high levels of hCD45^+^ cells in hOss, comprising MPO^+^/CD14^+^ granulo-monocytic cells, CD61^+^ megakaryocytes, glycophorin C^+^ erythroid cells and immature CD34^+^ HSPCs in F/hOss (**Figure 2B**). Qualitative and quantitative flow cytometry analysis further confirmed human multi-lineage hematopoietic formation in both F/hOss and P-N/hOss after 12 weeks HSPC post-transplant (**Figure 2C-I**). Erythroid cells were enhanced in F/hOss (10%) and to a lesser extent (albeit not significant) in P-N/hOss (6%) compared with mBM (2%) (**Figure 2D**), whereas hCD45^+^ cell percentages were similar in mBM from mice without and with hOss, regardless of the hMSCs origin (F/ or P-N/) (**Figure 2E)**. In terms of absolute numbers, a median of 2.8×10^6^ and 3×10^6^ hCD45^+^ cells/hOss were recovered from F/hOss and P-N/hOss at 12 weeks, respectively (**SupFig2B**). The most striking difference was the decreased percentage in CD19^+^ B-cells in F/hOss and P-N/hOss (46% and 53% of hCD45^+^ cells, respectively) compared with mBM from mice without hOss, in which B-cells predominated (80%) (**Figure 2F**) as typically observed in these humanized mouse models (Henry et al., 2020). Inversely, myeloid cell levels (CD14^+^/CD15^+^) were higher in hOss (34% and 27%) compared with mBM (10%) (**Figure 2G**). Only a minor difference was noted in B/myeloid/erythroid cell levels between F/hOss or P-N/hOss. Comparing B/myeloid/erythroid cell levels between mBM from mice with and without hOss showed a similar, albeit less pronounced, decrease in B-cell levels and an increase in myeloid cell levels in mBM from mice with hOss (**SupFig2E-H**), thus highlighting possible cross-talk between hOss and mBM sites. Analysis of immature human cell compartments indicated similar levels of CD34^+^ HSPCs in F/hOss, P-N/hOss and mBM without hOss at 12 weeks post-injection of CD34^+^ cells (**Figure 2H**). Assessing more immature hCD45^+^Lin^-^CD34^+^CD90^+^ HSCs indicated slightly enhanced levels of F/hOss (2.7%, p<0.05) compared with P-N/hOss (1.8%) and mBM (1.4%) (**Figure 2I**). Compared with mBM without hOss, CD34^+^ cells were significantly increased in mBM with F/hOss and mBM with P-N/hOss, while no differences were observed in hCD45^+^Lin^-^CD34^+^CD90^+^ HSCs (**SupFig2I-J**). These results show that hOss enable human hematopoiesis development, and differences were detected primarily in phenotypically mature hematopoietic cells compared with mBM. Interestingly, levels of B-cells, myeloid cells, and immature cells in hOss fell between the levels observed in mBM and hBM from young healthy donors (median age: 11 years) (**Figures 2F-I**), thereby suggesting that hOss better mimics normal human hematopoiesis.

**Figure 2.**
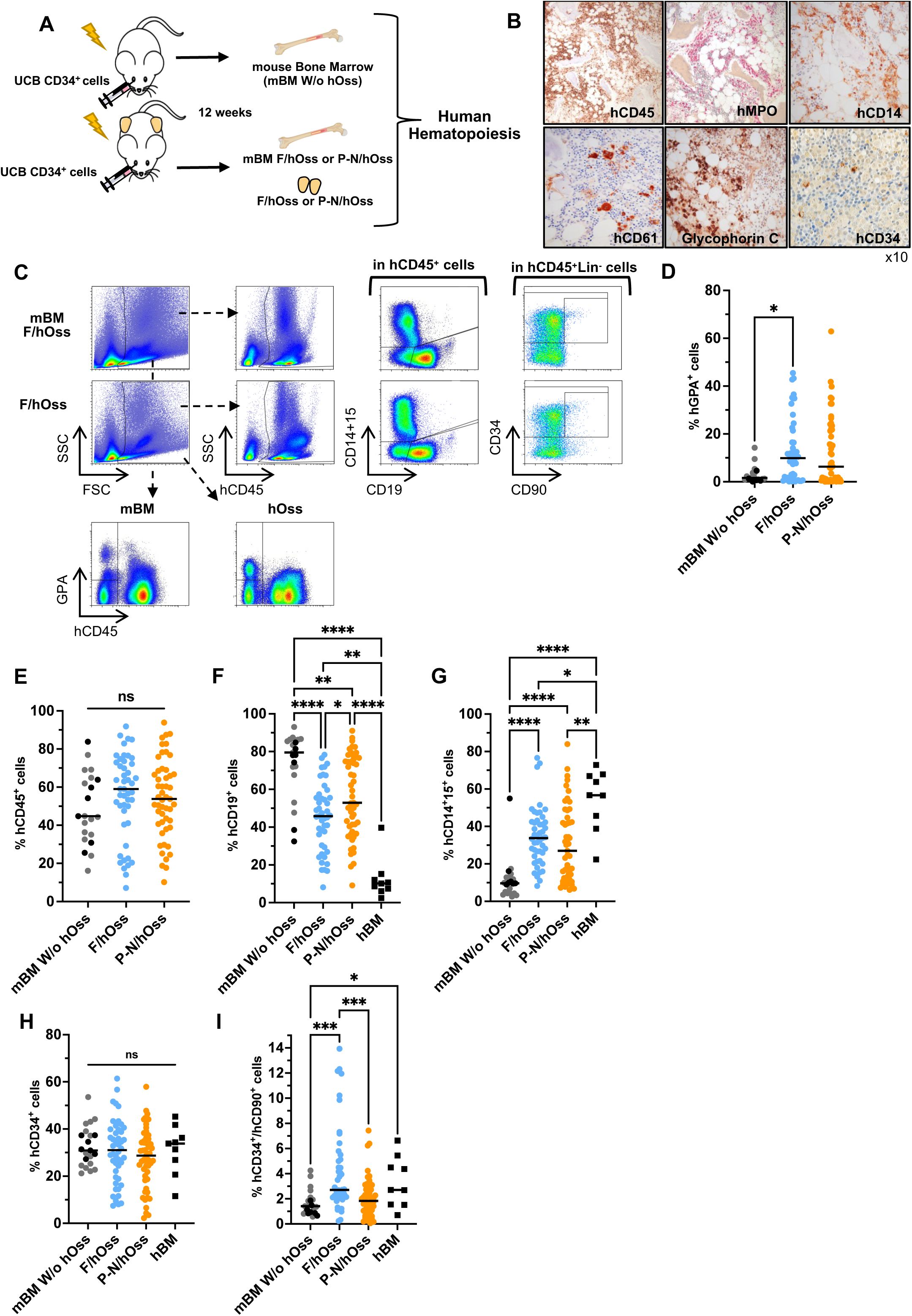
Human fetal hOss and post-natal hOss support normal human hematopoiesis. A. Experimental design of xenografts of umbilical cord blood CD34^+^ cells in mice carrying hOss. Umbilical cord blood CD34^+^ cells (10^5^ cells/mouse) were injected into mice carrying hOss and non-hOss controls, less than 24 hours after sublethal irradiation, 2-8 weeks after hOss implantation. Human hematopoiesis was analyzed 12 weeks after umbilical cord blood HSPC injection into the bone marrow of mice (mBM) and the hOss. B. Umbilical cord blood CD34^+^ injected in F/hOss at 12 weeks with histology immunolabeling of sections: human hematopoietic cells (hCD45), myeloid cells (hMPO, CD14), megakaryocytes (CD61), red cells (Glycophorin C), immature HSPCs (CD34). Amplification, ×10. C. A representative flow cytometry analysis of the human hematopoietic cells developing in mBM (upper panel) and F/hOss (lower panel). Analysis at 12 weeks post-injection of umbilical cord blood CD34^+^ cells. D. Percentage of human GPA^+^ cells in mice with and without hOss. Cells were gated on FSC/SSC following the gating strategy in C. Each dot represents individual mice (n=21 mice, without hOss in black and gray from 2 independent experiments; n=25 mice, with F/hOss, 47 hOss obtained from 5 different MSCs; n=28 mice with P-N/hOss, 51 hOss obtained from 4 different MSCs). Gray and black dots: mBM from mice without hOss; blue dots: F/hOss; orange dots: P-N/hOss. E. Percentage of cells expressing hCD45^+^ using the gating strategy shown in C for mBM (pooled 4 long bones) and hOss (2/mouse, treated separately) 12 weeks post-transplant of umbilical cord blood CD34 cells. The results represent the same mice as those displayed in D. F-G. Percentage of human (F) lymphoid CD19^+^ B-cells and (G) myeloid CD14^+^ /CD15 cells. Cells were gated on hCD45^+^ as described in C. hBM cells from healthy donors (age <18 years, n=9) were included as controls. The results represent the same mice as those displayed in D-E. The gating strategy is shown in SupFig 2D and Figure 2C. H-I. Percentage of CD34^+^ (H) and CD34^+^ CD90^+^ HSCs (I) gated on hCD45^+^ Lin (CD19 CD14 CD15) cells, based on the gating strategy described in C. hBM cells from healthy donors (age <18 years, n=9) were included as controls. The results represent the same mice as those displayed in D-E. ns, no statistical difference; *, p<0.05; **, p<0.001; ****, p<0.0001; Kruskal-Wallis test without correction.

### Fetal and post-natal hOss contain functional human mesenchymal stem cells

To characterize the possible replenishment of hOss by engrafted resident hMSCs, we tested whether isolation of hMSCs from primary hOss was possible and whether these recovered hMSCs could generate secondary hOss. After crushing primary F/hOss or P-N/hOss, adherent cells were recovered and amplified in plastic plates (**Figure 3A**). Flow cytometry analysis of adherent cells (using the same markers as displayed in **Figure 1B**) confirmed that the cells originated from human mesenchymal cells (**Figure 3B**). The secondary isolated hMSCs were injected into mouse flanks to form hOss, and human CD34^+^ cells (10^5^ cells/mouse) were transplanted 8 weeks later (**Figure 3A**). Upon sacrifice of the mice at 12 weeks post-transplant, secondary hOss had formed; interestingly, secondary F/hOss were twice as large (mg) as secondary P-N/hOss (**Figures 3C-D**). This difference was slightly less pronounced than that observed in primary hOss (**Figure 1E and SupFig2B**). Murine erythroid and hematopoietic cells were present in secondary hOss that did not receive CD34^+^ umbilical cord blood cells, although lower levels were observed compared with mBM especially in secondary P-N/hOss, within the limits of testing only a few hOss (**Figure 3E**). Human erythroid cells were enhanced in secondary hOss, as observed in primary hOss. However, P-N/hOss were less permissive to this human lineage development than F/hOss (**Figure 3F**). The percentage of hematopoietic cells did not vary between mBM and secondary F/hOss, but were significantly higher in secondary P-N/hOss compared with mBM (**Figure 3G).** In terms of absolute number of hCD45^+^ cells A significant decrease was observed in secondary P-N/hOss compared with F/hOss (**SupFig3A**). Compared with mBM from mice without hOss, secondary F/hOss displayed a significant decrease in CD19^+^ B-cells and a significant increase in CD14^+^/CD15^+^ myeloid/GPA^+^ erythroid cells, while a difference was not observed in secondary P-N/hOss (**Figure 3H-I**). Analysis of immature cells revealed similar CD34^+^ HSPC levels, and increased CD34^+^CD90^+^ HSC levels were observed only in secondary P-N/hOss compared with mBM (**Figure 3J-K**). As analyzed in mice with primary hOss, mBM from mice with secondary hOss displayed similar human hematopoietic reconstitution to that observed in secondary hOss, again indicating cross-talk between the mBM and hOss sites (**Figure 3F-K**). Analysis of hematopoietic cells from primary and secondary hOss further confirmed subtle but significant differences between F/hOss and P-N/hOss. Indeed, F/hOss appeared to effectively support balanced production of human lymphoid and myelo-erythroid cells, whereas secondary P-N/hOss displayed reduced development of human myelo-erythroid cells (**SupFig3B-E**). Furthermore, no major difference in the immature cell compartments was observed between primary and secondary hOss, although subtle but significant differences were observed in CD34^+^CD90^+^ cells (**SupFig3F-G**). Overall, these results show that hOss are dynamic human BM-like structures that contain hMSCs capable of serial production of hOss that continue to support human hematopoietic development. Interestingly, the lympho-myeloid bias observed in primary P-N/hOss was no longer observed in secondary P-N/hOss, thus suggesting that exhaustion may occur during the transplant process.

**Figure 3.**
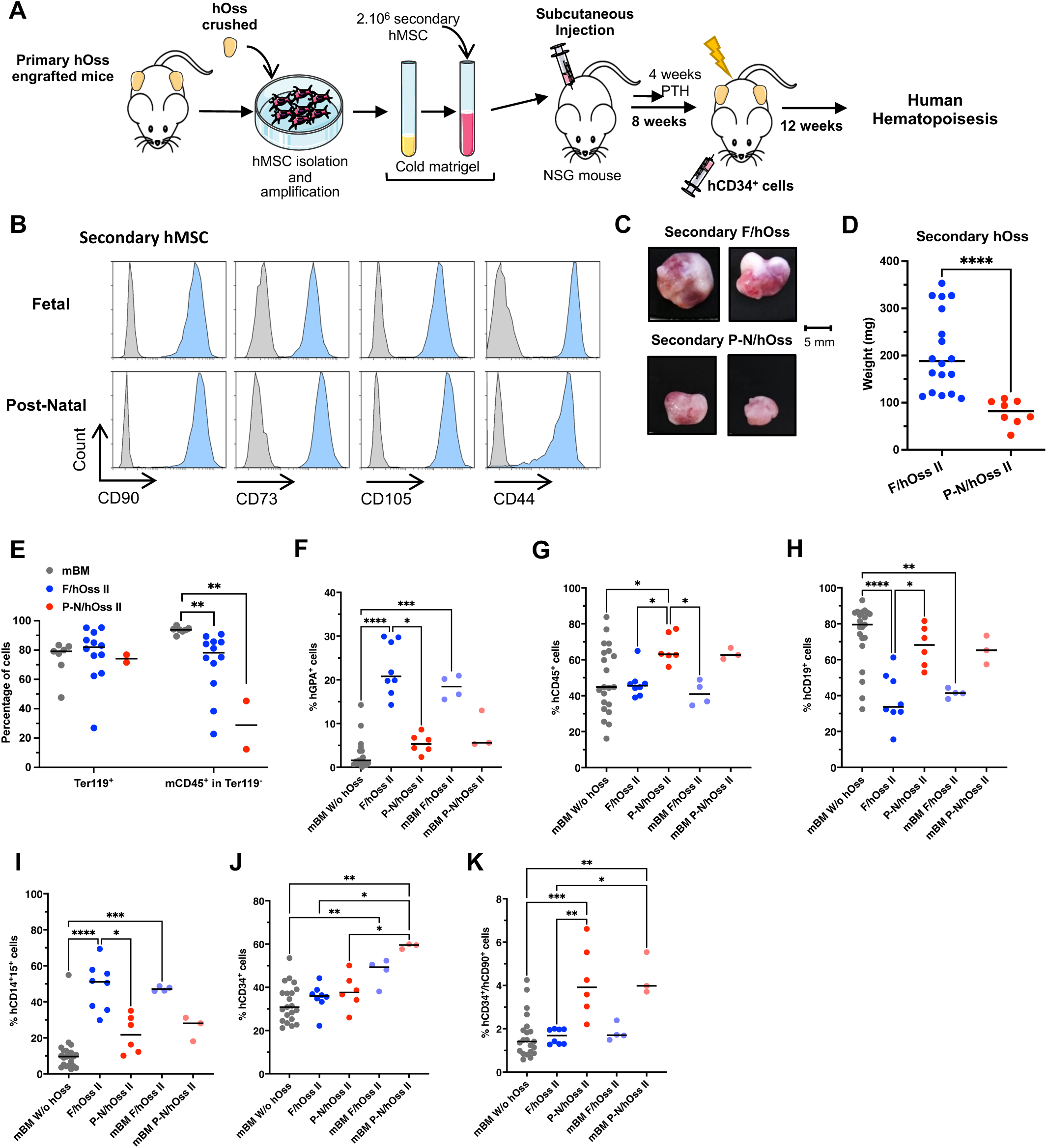
Human primary hOss comprise hMSCs capable of secondary hOss development. A. Experimental design of secondary hOss formation and grafts of CD34^+^ cells in mice carrying secondary hOss. Secondary hMSCs were isolated after crushing primary hOss, expanded as adherent cells and transplanted as indicated in Figure 1C. Eight weeks later, 10^5^ umbilical cord blood CD34^+^ cells were injected into mice carrying secondary hOss and non-hOss controls less than 24 hours after sublethal (2 Gy/mouse) irradiation. Human hematopoietic development was analyzed 12 weeks later in the bone marrow (BM) of mice and hOss. B. Flow cytometry analysis of secondary hMSCs isolated from primary F/ and P-N/hOss. Cells were gated on hCD45^-^ CD14^-^ CD15^-^ CD31^-^ cells (>90%). Representative of >4 tested secondary hMSC samples. The results were obtained from the gating strategy shown in SupFig1A. C. Representative examples of secondary hOss obtained 10-12 weeks post-injection of secondary hMSCs. The images display the results for the samples F28 (fetal hMSC origin, 11.6 PCWs) and ALLO3 (post-natal hMSC origin, 5 years old). D. Weight comparison of secondary F/hOss and P-N/hOss. The results were obtained from 18 hOss (individual blue dots, median value: black line) generated from secondary F/hMSCs, derived from 2 MSCs samples, in 3 independent experiments, and obtained from 8 hOss (individual red dots, median value: black line) generated from secondary P-N/hMSCs, derived from 1 hMSC sample in 2 independent experiments. E. Percentage of murine cells analyzed by flow cytometry, recovered from secondary hOss without injection of umbilical cord blood CD34^+^ cells. Gating was carried out on viable murine red blood cells, Ter119^+^, and viable murine hematopoietic cells (in Ter119^-^), mCD45^+^. The results are shown for 12 hOss generated with 1 secondary F/hMSC in 2 independent experiments and 2 hOss from 1 secondary P-N/hMSC in 1 experiment. F. Percentage of human GPA^+^ erythroid cells in mBM from mice with and without secondary (II) hOss. The gating strategy is the same as in Figure 2C. The results shown represent the individual values of mBM from 21 mice without hOss (gray dots: the same mice as displayed in Figure 2D), 8 hOss generated by 2 secondary F/hMSCs in 4 mice (dark blue dots), 6 hOss generated by 1 secondary P-N/hMSC in 3 mice (dark red dots), 6 mBM from mice with secondary F/hOss (light blue dots) and 3 mBM from the mice with secondary P-N/hOss (pink dots). Black lines indicate the median values. G. Percentage of hCD45^+^ cells assessed via flow cytometry in secondary hOss at 12 weeks post-transplant of umbilical cord blood CD34^+^ cells. The results represent the same mice as those displayed in F. The gating strategy is the same as that shown in Figure 2C. H-I. Percentage of human (H) lymphoid B-cells (CD19^+^) and (I) myeloid cells (CD14^+^/CD15^+^) gated on hCD45+ cells according to the strategy shown in Figure 2C. The results represent the same mice as those displayed in F-G. J-K. Percentage of (J) CD34^+^ and (K) CD34^+^ CD90^+^ HSCs gated on hCD45^+^ Lin^-^ (CD19^-^ CD14^-^ CD15^-^) cells, based on the same gating strategy as that shown in Figure 2C. The results represent the same mice as those displayed in F-G. *, p<0.05; **, p<0.01; ***, p<0.001; ****, p<0.0001; F-K. Kruskal-Wallis test without correction. D: Mann-Whitney test.

### Immature human hematopoietic cells from hOss display enhanced hematopoietic stem cell functional potential

We next characterized the immature cell function of hOss-engrafted CD34^+^ HSPCs using standard assays, such as myelo/erythroid colony forming units (CFUs) to measure committed progenitor levels and secondary reconstitution of immune-deficient mice to detect HSCs (Robin et al., 1999). Cells were obtained from primary hOss and the BM of mice with and without hOss as controls. Human CD34^+^ HSPCs were quantified and either plated in semi-solid methylcellulose medium or transplanted into NSG mice (**Figure 4A**). According to the CFU results, we observed similar total numbers of colonies generated by cells isolated from hOss and mBM (**Figure 4B**). Assessing colony types (i.e., progenitors) between the three conditions, we did not observe major differences in progenitor composition (**Figure 4C and SupFig4**). Analysis of the human hematopoietic cell levels in secondary NSG mice injected with cells from mBM of mice with and without hOss as well as primary and secondary hOss showed increased reconstitution potential of human HSPCs recovered from mice with hOss compared with mice without hOss, regardless of whether the hOss were primary or secondary (**Figure 4D-I**). Mice injected with cells from mBM with hOss also displayed hCD45^+^ cells (**Figure 4E and H)**, of which levels were greater than in mice with cells from mBM without hOss and similar or reduced compared with that observed in mice receiving cells from hOss (**Figure 4E and H)**. The human hematopoietic cells recovered from secondary NSG mice were primarily composed of B lymphoid cells, regardless of origin (hOss and mBM) (**Figure 4F and I**). Overall, these results show that hOss maintain stronger human HSC potential than mBM of mice without hOss. Interestingly, the enhanced myeloid cell differentiation observed in hOss (**Figures 2 and 3**) was lost in the mBM of secondary NSG mice in the absence of the hOss environment.

**Figure 4.**
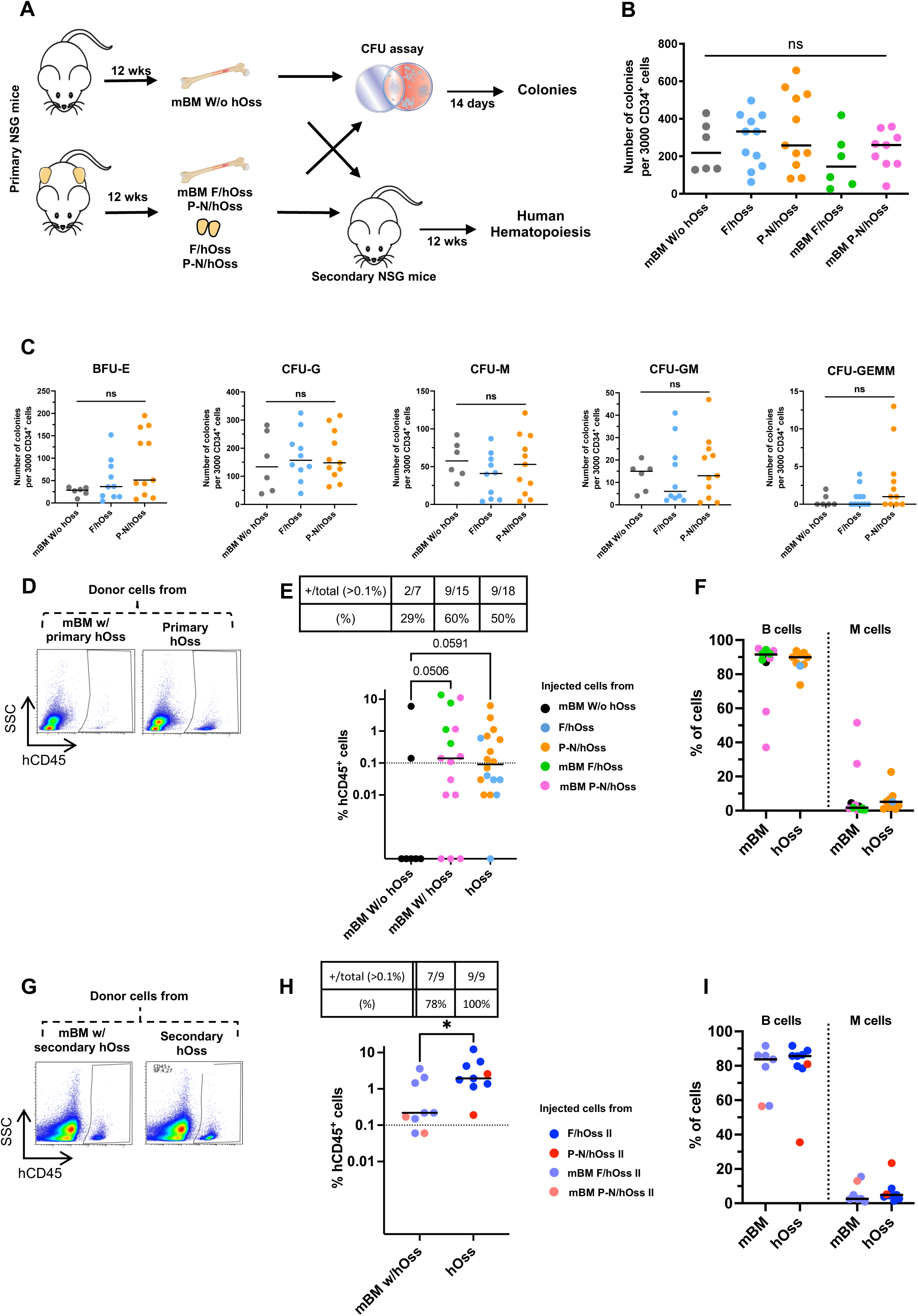
Human primary hOss display enhanced levels of functional immature human cells. A. Experimental design of the functional analysis of the immature cell compartment in primary hOss compared with murine bone marrow (mBM) from mice with and without hOss. CD34^+^ cells recovered from crushed hOss and mBM 12 weeks after CD34^+^ cell transplant were analyzed by flow cytometry and either plated in methylcellulose semi-solid medium (1×10^3^ hCD34^+^ cells/plate) or re-injected into secondary NSG mice (3-5.10^5^ hCD34^+^cells/mouse). Two weeks later, colonies were scored and characterized as granulocytic (CFU-G), monocytic (CFU-M), granulo-monocytic (CFU-GM), erythroid (BFU-E) or multipotent (CFU-GEMM). Human hematopoiesis was analyzed in the BM of secondary mouse recipients 12 weeks post-injection. B. Total number of colonies generated by 3,000 plated CD34^+^ cells at 2 weeks of culture, isolated from different BM sites (mBM without hOss, gray dots; F/hOss, blue dots; P-N/hOss, orange dots; mBM with F/hOss, green dots; mBM with P-N/hOss, pink dots). The results were obtained from 3 independent experiments and each sample was tested in triplicate. The individual mean of triplicates for each tested sample is shown; black lines indicate the median values. ns, not significant; Kruskal-Wallis test without correction. C. Total number of colonies per colony type generated by 3,000 plated CD34^+^ cells at 2 weeks of semi-solid cultures (mBM with hOss, gray dots; F/hOss, blue dots; and P-N/hOss, orange dots). Black lines indicate the median values. ns, not significant; Kruskal-Wallis test without correction. D. Representative examples of FACS plots of hCD45^+^ cells detected in the mBM of NSG mice injected with cells from mBM of mice with hOss (primary) and hOss (primary). The cells were gated on FSC/SSC parameters. E. Upper panel: ratio of secondary mice engrafted with >0.1% of hCD45+ cells on all injected mice. The associated proportion was calculated. Lower panel: percentage of hCD45 cells analyzed in the BM of NSG mice injected with cells from mBM mice without (7 mice) and with primary hOss (15 mice) and from primary hOss (18 mice). Black lines indicate the median values. Kruskal-Wallis test without correction. F. Relative percentage of B lymphoid (CD19^+^) and myeloid (CD14^+^/CD15^+^) cells from the pooled mice shown in E. Only the mice with reliable engraftment (≥0.1% hCD45 cells) were analyzed. Black lines indicate the median values. G. Representative examples of FACS plots of hCD45^+^ cells detected in the mBM of NSG mice injected with cells from mBM of mice with hOss (secondary) and hOss (secondary). The cells were gated on FSC/SSC parameters. H. Upper panel: ratio of secondary mice engrafted with >0.1% of hCD45^+^ cells on all injected mice. The associated proportion was calculated. Lower panel: percentage of hCD45 cells analyzed in the BM of NSG mice injected with cells from secondary hOss (9 mice,) and mBM of the same mice (with hOss, 9 mice). Black lines indicate the median values. *, p<0.05; Mann-Whitney test. F. Relative percentage of B-lymphoid (CD19^+^) and myeloid (CD14^+^/CD15^+^) cells from the pooled mice shown in H. Only the mice with reliable engraftment (≥0.1% hCD45^+^ cells) were analyzed. Black lines indicate the median values.

### Gene expression profiles reveal enhanced myeloid precursors/progenitors and decreased lymphoid precursors/progenitors in immature human hematopoietic cells from fetal and post-natal hOss

To further characterize the immature cell compartment engrafting hOss compared with mBM from mice without hOss, we performed single-cell RNA sequencing of hCD34^+^ cells sorted at 12 weeks post-transplant (**Figure 5A**). A total of 9,894 cells, containing 3,188, 4,005 and 2,701 cells from mBM without hOss, F/hOss and P-N/hOss, respectively, were successfully integrated in UMAP (**SupFig5A-B**). Seurat unsupervised clustering (resolution levels = 0.2) was applied to separate cells into 13 clusters according to the gene expression profile (Hao et al., 2021). Comparing these results with the expression profiles described in (Hay et al., 2018), the 13 clusters were classified according to cell differentiation pathway, which highlighted lymphoid (CD34^+^ Multilin/CLP, CD34^+^ pre-B cycling, CD34^+^ pro-B cycling, CD34^+^ pre-plasma cell (Pre-PC)), myeloid (immature neutrophils/monocytes, neutrophils), erythroid and dendritic cell populations (CD34^+^ early erythroblasts, erythroblasts, CD34^+^ MDP/pre-dendritic and dendritic) (**Figures 5B-C**). Cell cluster annotation revealed that 70.5% of cells (6,975 cells) were either B-cell lineage oriented (CD34^+^ pre-PC and pre-B cell cycling) or multilineage HSCs/progenitors (CD34^+^ Multilin/CLP and CD34^+^ HSC/MPP/LMPP) (**Figure 5D**). The other immature cells (immature neutrophils/monocytes, CD34^+^ MDP/pre-dendritic cells, CD34^+^ Eo-B-Mast, CD34^+^ early erythroblasts, erythroblasts) were detected at lower levels. The repartition of these cell subpopulations/clusters indicated variations among cells isolated from hOss and mBM; for example, early erythroblasts and neutrophils (clusters 7 and 9) were frequent in P-N/hOss cells, while CD34^+^ eosinophils-basophiles-mastocytes (cluster 8) were highly represented in F/hOss (**SupFig5C).** When progenitors/precursors were grouped into B lymphoid (including CD34^+^ pre-PC, CD34^+^ pre-B cycling, CD34^+^ Multilin/CLP, CD34^+^ pro-B, CD34^+^ pre-B, Follicular B cells), myeloid/erythroid (immature neutrophils/monocytes, neutrophils, CD34^+^ MDP/pre-dendritic cells, CD34^+^ Eo-B-Mast, CD34^+^ early erythroblasts, erythroblasts) and early HSPC (CD34^+^HSC/MPP/LMPP) lineages, significant differences were observed between mBM and hOss (**Figure 5E**) that were similar to variations observed in the mature cell compartments (**Figure 2E-I**). Indeed, this analysis showed that immature cells from F/hOss and P-N/hOss comprise significantly reduced B lymphoid precursors/progenitors (p<0.0001, χ^2^ test) and increased myeloid/erythroid precursors/progenitors (F/hOss *vs* mBM, p<0.0001; P-N/hOss *vs* mBM, p<0.0001, χ^2^ test) compared with mBM from mice without hOss. We also observed a slightly enhanced proportion of HSC/MPP/LMPP cells in hOss (**Figure 5E**, F/hOss *vs* mBM, p<0.0001, P-N/hOss *vs* mBM, p=0.0015, χ^2^ test), in accordance with the results of the functional assays displayed in **Figure 4C-F**.

**Figure 5.**
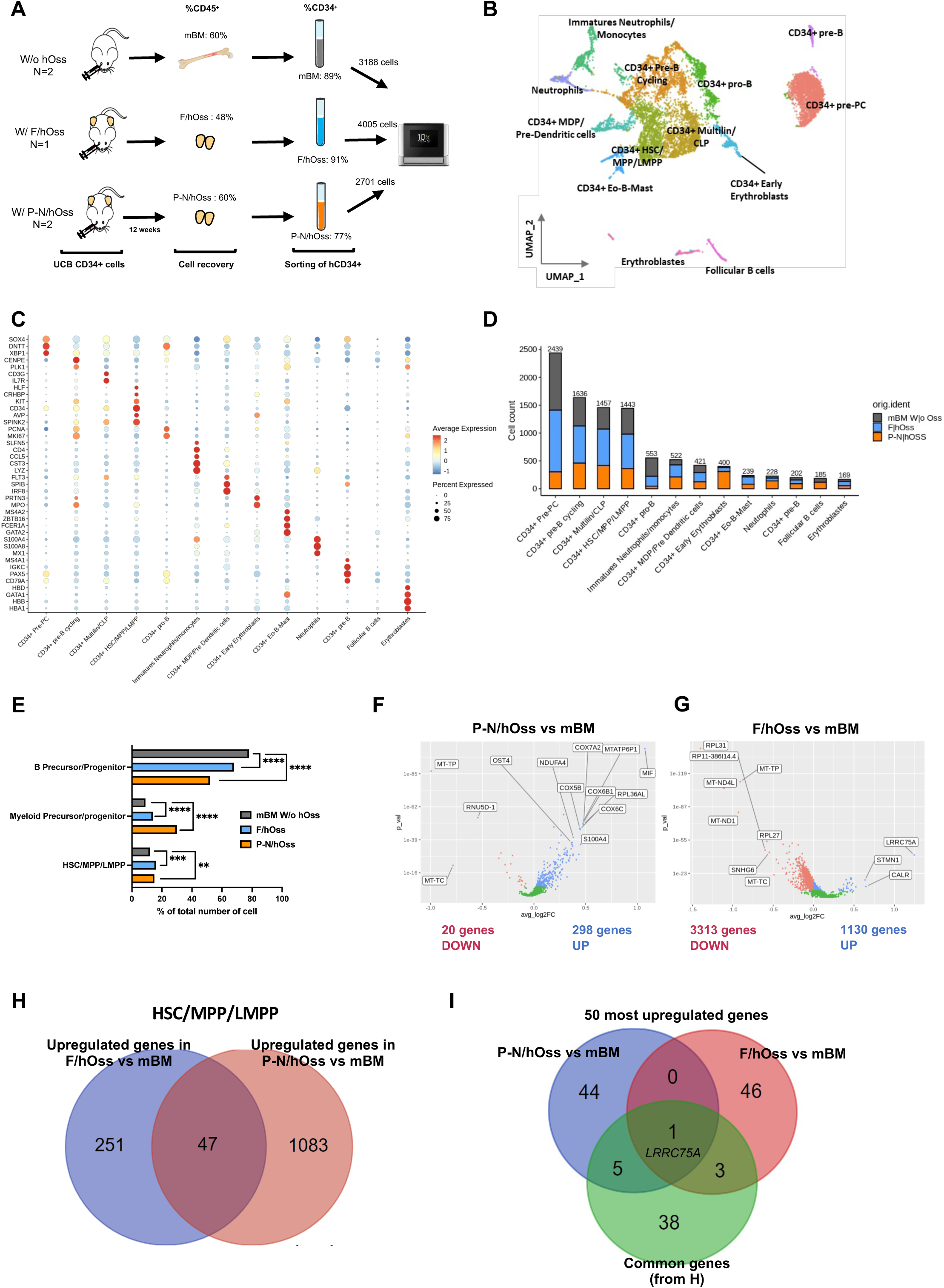
Single cell analysis shows that immature cells recovered from hOss display an enhanced bias toward myelo/erythroid progenitors and increased HSC levels. A. Experimental design. Three months after injection of 10^5^ umbilical cord blood CD34^+^ cells, mononuclear cells were isolated from mBM and hOSS and pooled according to origin. CD34^+^ cells were then sorted with a purity >70% and viability >90%. Sorted cells were then used to prepare single-cell RNA-sequencing libraries. B. UMAP representation of the 13 clusters identified via Seurat. Each cluster is represented by a different color. Clusters were annotated by comparing the gene expression profiles with those of the hematopoietic populations described by Hay et al. (Hay et al., 2018a) (HSC = hematopoietic stem cell, MPP = multipotent progenitor, LMPP = lymphoid-primed multi-potential progenitor, MDP = monocyte/dendritic cell progenitor, CLP = common lymphoid progenitor, Multi-Lin = multi-lineage progenitor, pre-PC = pre-plasma cells, Eo-B-Mast = eosino-baso-mast cells). C. Annotation of the 13 clusters according to gene markers as previously described (Hay et al., 2018b; Baccin et al., 2020). The color code is shown in the figure legend. D. Comparison of population distribution (absolute numbers) between the mBM and hOss compartments. Each bar corresponds to the total number of cells in a given sub-population. The sum of each bar corresponds to the total number of cells in our dataset. Each bar is divided into 3 segments corresponding to the original samples. E. Comparison of the proportion of 3 major progenitor groups (HSC/MPP/LMPP, B cell precursors/progenitors and myeloid precursor/progenitor cells) in the mBM and hOss conditions. Proportions were calculated within each sample using the data from E. “B Precursor/Progenitor” include CD34^+^ pre-PC, CD34^+^ pre-B cycling, CD34^+^ Multilin/CLP, CD34^+^ pro-B, CD34^+^ pre-B, follicular B cells. “Myeloid precursor/progenitor” contain immature neutrophils/monocytes, neutrophils, CD34^+^ MDP/pre-dendritic cells, CD34^+^ Eo-B-Mast, CD34^+^ early erythroblasts, erythroblasts. Early HSPC are contained in CD34^+^ HSC/MPP/LMPP cells. The proportions were compared using a χ2 test. For each test, **p < 0.01, ***p < 0.001, ****p < 0.001. F-G. Differentially expressed genes between post-natal hOss (F) or fetal hOss (G) and mBM cells within the CD34^+^ HSC/MPP/LMPP compartment. Each volcanoplot indicates the expression of genes that were significantly downregulated (p < 0.05, red dots) or upregulated (blue dots) in hOss compared with mBM. Genes that were not significantly underexpressed or overexpressed are shown in green. H. Venn Diagram showing the 47 common upregulated genes in the CD34^+^ HSC/MPP/LMPP compartment from P-N/hOss (blue) and F/hOss (red), both compared with mBM. I. Venn diagram showing 9 out of the 47 gene list (green) from H identified among the 50 most upregulated genes, P-N/hOss compared with mBM (blue), and F-hOss compared with mBM (red).

We also assessed whether hOss could impact gene expression in the most immature CD34^+^ HSC/MPP/LMPP cells of our data set. We analyzed the most differentially expressed genes in the HSC/MPP/LMPP cell cluster isolated from hOss and mBM. Comparing P-N/hOss and mBM, we identified 318 differentially expressed genes (p<0.05), of which 298 genes were upregulated and 20 genes were downregulated in P-N/hOss-derived cells (**Figure 5F**). Comparing F/hOss and mBM, we found 4,443 differential genes (1,130 upregulated genes and 3,313 downregulated genes in F/hOss) (**Figure 5G**). The 50 most upregulated genes are presented in **SupTables 1 and 2**. Examination of the two lists of upregulated genes revealed 47 common genes, of which 9 genes were included among the 50 most upregulated genes in both F/hOss and P-N/hOss (**Figure 5H-I, SupTable 3, SupTable 1-2**). Among these 9 genes, *LRRC75A* (Leucin-rich Repeat Containing 75A protein) has recently been implicated in the regulation of VEGF in ischemic BM-MSC, and VEGF is a regulator of HSC function (Miura et al., 2023). However, *VEGF* was not enhanced in our HSC/MPP/LMPP data set, thus suggesting that *LRRC75A* enhancement does not impact *VEGF* transcriptional levels in immature cells, in accordance with the finding that VEGF is typically produced by endothelial cells. To further examine the modified processes in hOss-derived cells, we performed KEGG and GO enrichment analyses using the 100 most deregulated genes in P-N/hOss and F/hOss, compared with mBM. We found that metabolic pathways/OXPHOS-cell respiration was enriched in P-N/hOss, and several pathways unrelated to HSC function/potential were slightly enriched in F/hOss (**SupFig5B-C**). We also assessed several other gene sets more specific to hematopoietic cells (list in **SupTable4**) but did not observe enrichment in any specific pathway in HSC/MPP/LMPP cells from hOss compared with mBM. Overall, these results revealed only a discreet qualitative distinction between the HSC compartments recovered from hOss and mBM. However, we observed a slightly increased proportion of HSCs in hOss and a bias toward B lymphoid/myelo-erythroid differentiation at the progenitor/precursor levels, which may contribute to the observed mature cell variations and the enhanced functional HSC.

### Clonal tracking shows that hOss promote the human myelopoietic cell lineage in both hOss and the mBM of mice with hOss

We next used lentiviral barcoding to label and track CD34^+^ HSPC-driven hematopoietic reconstitution at the clonal level in hOss and mBM (**Figure 6A**). CD34^+^ cells from umbilical cord blood were transduced with 21-bp lentiviral barcode libraries containing 40×10^4^ different barcodes expressing GFP (Eisele et al., 2022). Low transduction efficiency was achieved (7.8-16.8% GFP^+^ cells, **SupFig6A and SupTable5)** ensuring that most cells were labeled with one barcode. Transduced cells were injected into mice with or without hOss and hematopoietic GFP^+^ CD19^+^ B-cells, CD14^+^CD15^+^ myeloid cells and CD34^+^ immature cells were sorted from hOss and mBM at 12 weeks after injection (**Figure 6A, SupFig6B**). At that time point, GFP^+^ and GFP^-^HSPCs had generated the same proportion of each cell lineage fraction (**SupFig6C**), thus showing similar differentiation from the transduced and non-transduced cells. After filtering and quality control (**SupFig6D**), we obtained variable barcode numbers between mice, which was neither correlated with the presence of hOss nor the hOss origin (**SupTable5**). Based on the number of injected CD34^+^ umbilical cord blood cells and the transduction efficiency, we estimated the frequency of HSCs at 1/1000 to 1/100 (**SupTable5**), which is in accordance with a previous study using barcoded human HSCs (Cheung et al., 2013). We then assessed hematopoietic variations between mice with and without hOss. In the mBM without hOss, the progeny of barcoded HSPCs was primarily biased toward B-cells, whereas in mice with hOss, the barcoded HSPCs more uniformly yielded B-cells and myeloid cells (**Figure 6C),** which is in agreement with the flow cytometry results (**Figure 2**). In mice bearing hOss, we found an increased proportion of clones containing myeloid cells, which was only statistically significant for mice with P-N/hOss (**SupFig6E**). No difference was observed in total barcodes (**Figure 6B)** or the number of cells produced per barcode (clone size) when hOss and mBM of mice with hOss were compared with mBM without hOss, with the exception of myeloid clones in the P-N/hOss condition, which were significantly smaller (**SupFig6F)**. Thus, the increased myeloid to B lymphoid ratio observed at the population level in P-N/hOss was associated with an increase in small myeloid-biased clones and not to an increased production of total myeloid cells per clone. Next, we investigated the potential relationship between human hematopoiesis in hOss and mBM in the same mice. More than 40% of clones/barcodes were common between the hOss and mBM (**Figures 6D-E**, **blue boxes, SupFig6G**), thus indicating that many hematopoietic cells originating from the same HSPC were distributed in both sites. These observations provide insight into the mechanisms of phenotypic similarities between mBM from mice with hOss and hOss-containing cells at the population level (**SupFigures 2C-F**). The common cells constituted mainly large multipotent clones in which CD34^+^ immature cells were also detected, which is indicative of self-renewal properties (**Figures 6D-E**). Clones uniquely present in one BM site were also detected (**Figures 6D-E, red boxes, SupFig6G**) and found at similar levels in mBM and F/hOss, whereas more unique clones were detected in P-N/hOss (**SupFig6G)**. Interestingly, while mice with F/hOss were primarily reconstituted with mBM/hOss common large multipotent clones (**Figure 6D, SupFig6H**), mice with P-N/hOss displayed clones that varied in size, with a trend of enhanced myeloid-restricted clones that were either common or localized in P-N/hOss (**Figure 6E**, **SupFig6I)**. Overall, our results indicate that a diversity of multipotent or more lineage-restricted HSPC clones reconstitute human hematopoiesis in both the mBM and hOss in these models, many of which were common between BM sites. The presence of hOss creates a more favorable environment for human myelopoiesis in large clones in F/hOss and small clones in P-N/hOss.

**Figure 6.**
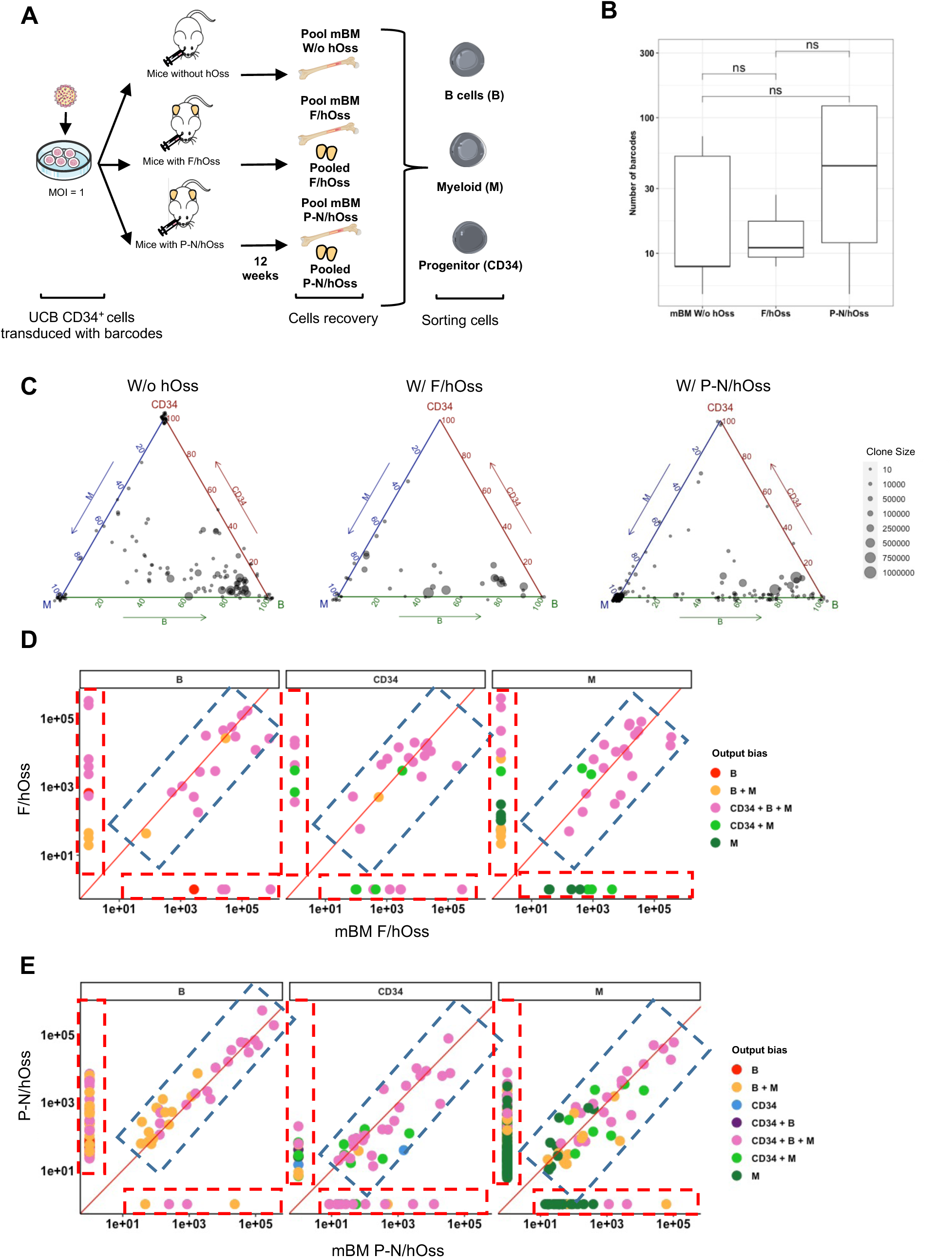
Cellular barcoding shows that hOss support more balanced human hematopoietic development. A. Experimental design. Injection of 10^5^ umbilical cord blood CD34^+^ cells was done after barcode-transduction. Three months later, mononucleated cells were isolated from mBM and hOss and pooled according to origin. Transduced (GFP+) B, Myeloid (M) and CD34^+^ cells were then sorted, lyzed and barcodes were amplified by PCR for identification and analysis. B. Number of unique barcodes identified per mouse in each condition. ns, not significant; t-test. C. Ternary plot of the fraction of cells produced per barcode clone in each of the 3 cell types (CD34^+^, B-cells, and myeloid (M) cells). Each dot represents a distinct barcode. The size of the dot indicates the number of cells originating from each barcoded progenitor injected (clone size). D-E. Number of reads per barcode clones detected in the hOss and bone marrow for F/hOss (D) and P-N/hOss (E) in the various cell types. Each dot represents a distinct barcode. The red line represents the point where barcodes would have the same clone size in both organs. The blue boxes contain the common barcodes/clones and the red boxes contain barcodes/clones specific to mBM and hOss. The axis is log10 transformed of the renormalized read count+1. The threshold for lineage content is 1%, and the minimal clone size is 10 cells.

## Discussion

In this study, we show that hBM hOss models support the capacity of human hematopoietic cell differentiation and production in immune-deficient mice harboring hOss. These hOss also exhibit intrinsic self-replenishing potential by maintaining implanted hMSCs capable of serial functional hBM formation and hematopoietic support. These results are concordant with previously described hBM models that display efficient human hematopoietic support following HSPC transplant, including balanced production of mature myeloid cells compared with mature lymphoid cells (Chen et al., 2012; Abarrategi et al., 2017; Fritsch et al., 2018; Grigoryan et al., 2022), thus demonstrating that the results between different hOss models are reproducible. Such a myeloid bias was detected, regardless of whether the MSCs used to form hOss were of fetal or P-N origin. However, P-N/MSC-derived hOss and mBM displayed smaller myeloid-biased HSPC clones compared with mice with F/hOss, in which myeloid cells were produced from more multipotent clones. These results suggest that even if the developmental origin of MSCs did not significantly impact hematopoietic cell type recovery at the population level, there are subtle differences between F/hOss and P-N/hOss niches in terms of the type of clones that reconstitute human hematopoietic cells.

It was previously unknown whether this bias in lineage cell production originated from biased differentiation of HSCs or progenitors towards the myeloid lineage at the expense of the lymphoid lineage. In our study, we assessed gene expression profiles in CD34^+^ cells isolated from hOss and mBM at the single-cell level. Our results indicate that HSPCs are not lineage bias at the transcriptional level, as specific lineage commitment programs were not observed in HSPCs isolated from hOss, compared with mBM. The single-cell RNA sequencing results rather highlighted the presence of larger myeloid and erythroid progenitor/precursor compartments, which is probably supported by human factors produced by the humanized microenvironment of hOss (Chen et al., 2012; Reinisch et al., 2016; Grigoryan et al., 2022). These results are in accordance with previous findings from a BM model generated by implanting collagen scaffolds carrying human MSCs, which displayed enhanced myeloid progenitors, as assessed by cell surface marker and CFU-C analysis (Fritsch et al., 2018). However, when secondary transplants were performed in regular NSG mice (without hOss), even though HSC levels were enhanced in both models (our results and Fritsch et al., 2018), we did not observe an increased myeloid bias in human CD45^+^ cells derived from hOss, thus indicating that without the hOss influence, the myeloid potential of human HSPCs is no longer detected. The weak enhancement in myeloid production observed in previous studies was probably due to the recipient mouse models used (STRG, MSTRG), which produce human myeloid cytokines (Fritsch et al., 2018). Our results indicate that mice with hOss can substitute cytokine-humanized mice in terms of myeloid cell lineage production from engrafted HSCs.

Indeed, using NSG recipient mice that do not produce human factors, we found that primary hOss had a strong impact on human hematopoiesis, which was detected from mature cells to immature cells, and was observed from hOss to the mBM of mice with hOss. Importantly, this impact on lineage properties persisted in MSCs isolated from primary hOss, as observed in human hematopoietic cells recovered from secondary hOss, especially when hOss formation was of F/MSC origin, thus suggesting that F/MSCs are more potent than P-N/hOss in the long term. These results indicate that hOss are dynamic multicellular structures that maintain MSCs with the ability to differentiate in hOss capable of perpetuating cell lineage in a balanced manner, which is an established feature of BM. Interestingly, in one study in which hOss were implanted after HSPCs had engrafted mBM, the balanced myeloid-lymphoid progeny was observed in hOss (Reinisch et al., 2015). This finding is in line with our results of mBM from hOss-bearing mice, thus displaying the impact of hOss both in proximity and at a distance. Using clonal tracking of hematopoietic cell production, we show that in hOss-grafted mice, many human hematopoietic cells recovered from mBM and hOss originated from multipotent HSPCs common between mBM and hOss, especially in F/hOss. Such clones were also detected independently in hOss or mBM, although they were rarer, thus highlighting that at 12 weeks post-transplant, human hematopoiesis in such models occurs through differentiation of multipotential HSCs. Furthermore, smaller unipotent myeloid cell clones were present in hOss, especially in P-N/hOss, potentially associated with different waves of progenitor differentiation or concomitant production of hematopoietic cells by a diversity of HSPCs, as previously described (Cheung et al., 2013). The extent to which hOss and mBM compete to attract HSCs versus committed progenitors remains to be assessed using hematopoiesis kinetic analyses. Whether the common clonally-related hematopoietic cells are due to cross-talk of differentiated cells between mBM and hOss or are rather generated from single barcoded HSCs that divided, trafficked through the blood stream and settled in the mBM and hOss to differentiate remains to be examined.

In this study, we also investigated whether the developmental age of BM from which MSCs were recovered impacted hOss formation and hematopoietic support. Previous studies on human F/MSCs, pediatric MSCs and adult MSCs have reported variations in cell surface marker expression and fibroblast CFUs, outlining specificities in F/MSCs (Maijenburg et al., 2012). In mice, several age-related BM microenvironment factors modulate HSC potency, including osteopontin, which enhances levels in young adults compared with older adults - attenuating aging of old HSCs, and the CCL5/RANTES, which is a secreted matrix protein that contributes to the bias in old HSCs toward myeloid differentiation (Guidi et al., 2017; Ergen et al., 2012). In our experiments, both F/MSC and P-N/MSC reproducibly formed hOss, with increased size and weight in F/MSC-derived hOss, which was likely associated with enhanced cell proliferation and thicker bone structure. Such differences did not dramatically impact hematopoietic cell production at the population level, at least in primary hOss, with a trend towards myeloid progenitors only detected by single-cell RNAseq in P-N/MSC-derived hOss compared with F/MSC-derived hOss. Interestingly, secondary hOss perpetuated the variations in size albeit to a lesser extent, and P-N/hOss lost the myeloid-lymphoid lineage balance, indicative of exhaustion most likely mediated by stress-induced serial transplantation. Further research is necessary to understand the drift mechanisms in primary MSCs undergoing hOss formation. The possible aging caveat of serial tissue culture-isolated MSCs and the difficulty to reproduce hOss formation from one lab to the next has recently been challenged (Côme et al., 2020). A human telomerase-immortalized mesenchymal cell line, Mesenchymal Sword of Damocles, has been shown to recapitulate the various multipotent and hematopoietic supportive properties of primary hMSCs by generating hOss in vivo after ex-vivo priming to chondrocyte or osteocyte differentiation (Grigoryan et al., 2022). Such a cell line is a valuable tool as it produces >1000 hOss originating from a clonal cell population (Grigoryan et al., 2022). hOss obtained from Mesenchymal Sword of Damocles cells and primary MSC samples are also useful recipients of pathologic cells, such as acute myeloblastic leukemia and BM-infiltrating solid tumor cells (breast cancer and neuroblastoma). This approach enables analysis of the relationship between the BM microenvironment and pathologic cells in a humanized context, including the soluble factors involved (Antonelli et al., 2016; Abarrategi et al., 2017; Reinisch et al., 2016; Vaiselbuh et al., 2010; Martine et al., 2017; Grigoryan et al., 2022).

These results contribute novel findings regarding the distinct impact of hBM models generated from hMSCs isolated from normal F and P-N donors on normal hematopoietic development after transplantation of human HSPCs. Continued advancements in these models, such as the incorporation of human endothelial cell-derived vasculature (Passaro et al., 2017), will undoubtedly further impact normal and abnormal hematopoietic cell development. However, detailed characterization at the clonal levels, as described here, is essential to validate the physiological relevance of these models to study normal and abnormal human cells.

## Materials and methods

### Human samples

Umbilical cord blood (CB) samples were collected from healthy infants with the informed written consent of the mothers based on the declaration of Helsinki. Samples were obtained in collaboration with Clinique des Noriets, Vitry-sur-Seine and the Cell Therapy Department in Hôpital Saint Louis, Paris, France. Human BM used in Figure 2F-J were collected from allografts (mostly children sibling donors) at the Cell Therapy Department in Hôpital Saint Louis, Paris, France. Samplings and experiments were acknowledged by the Institutional Review Board of INSERM (Opinion number 13-105-1, IRB00003888). CB and BM cells were subjected to Ficoll gradient. CB mononucleated cells were enriched for CD34^+^ cells using the human CD34 MicroBeads Kit following manufacturer instructions (130-046-703, Miltenyi Biotec), assessed for purity (≥70% CD34^+^ cells), and used directly in experiments or froze in fetal bovine serum (FBS, F9665, Sigma-Aldrich) supplemented with 10% DMSO (D5879, Sigma-Aldrich) for later usage. hBM cells were processed for Flow cytometry directly after Ficoll.

Fetal bone marrow (F/BM) samples to 12 weeks post-conception were obtained after abortion processes with the informed written consent of the parents in accordance to guidelines approved by the French Agence de Biomédecine (PFS18-009). Samples were obtained in collaboration with the department of Foeto-anatomo-pathology of the Hospital Antoine Beclère, Clamart, France. Post-natal BM (P-N/BM) MSC samples were collected from healthy children and adult allografts at Necker Enfants Malades hospital, France.

### Human MSC (hMSC) isolation and culture

Human F/BM samples were crushed in DPBS with 1% of penicillin-streptomycin (PS). Following centrifugation, the BM fragments were resuspended in α-MEM medium (M4526, Sigma-Aldrich) containing 0.22μm-filtered 10% human platelet lysate provided by the Department for Transfusion Medicine, Paracelsus Medical University, Salzburg, Austria or purchased from StemCell^TM^ Technologies (Stem Cells Lysate, 06962), 1% PS and glutamine, 2 units/mL heparin (hereafter called MSC medium) and incubated in plastic dishes. P-N/BM cells were extracted from filters, centrifuged and directly plated in MSC medium without Ficoll process. After amplification, adherent cells were split for further expansion (passage 0) or directly plated for experiments. At every passage, the culture medium was gently removed, plastic dishes were carefully washed with DPBS followed by addition of trypsin/EDTA solution (0.05%, 25300-054, Gibco) and incubated at 37°C for 5-10 minutes. Once hMSC were detached, αMEM medium containing 10% of FBS was added. Recovered adherent cells were resuspended in MSC medium after centrifugation and counted using the GuavaEasycyte 8HT apparatus in Muse Count & viability kit (MCH600103, Cytek Biosciences) or with trypan blue (T8154, Sigma-Aldrich). Part of cells were frozen at 1×10^6^ cells/vial in FBS containing 10% DMSO. Adherent cells were characterized by flow cytometry to measure their MSC content. To expand hMSC in cultures, a minimal number of 500 cells per cm2 were set at start. Adherent cells were amplified during maximal 4 passages in MSC medium. Secondary hMSC, isolated from primary hOss, were obtained using the same method as for primary hMSC isolation, e.g. after crushing of primary hOss, adhesion of cells on plastic dishes, expansion in hMSC medium and immunophenotyping to ensure their hMSC content.

To measure cell proliferation, hMSC were seeded at 500 cells/cm2 in 24-well plates in quadruplicates, three-days prior the counting was started. Thereafter adherent cells were detached every 24 hours and counted using the GuavaEasycyte 8HT with Muse Count & viability kit.

To test for adipocyte, osteocyte and chondrocyte differentiation, hMSC were cultured in conditions provided by Miltenyi Biotec using StemMACSTM Adipodiff media Human (130-091-677), StemMACS^TM^ OsteoDiff media Human (130-091-678) and StemMACS^TM^ ChondroDiff media Human (130-091-679). Oil Red staining kit (0843-SC, Cliniscience) was used to reveal adipocyte differentiation, and Alizarin Red S Staining kit (0223-SC, Cliniscience) helped showing osteocyte differentiation. All imaging were performed with the EVOS FL Auto Imaging System (Thermo Fisher Scientific).

### hMSC flow cytometry antibody panel

All antibodies were used at the 1/100 dilution except for CD44 labelling (1/500). Detached adherent cells were labelled with antibodies directed against human CD14 (clone 61D3), CD44 (IM7, both from Invitrogen) CD15 (W6D3), CD31 (WMD59), CD105 (SN6H), CD73 (AD2), CD45 (HI30, all from BioLegend®), CD146 (541-0B2, Miltenyi Biotech), and CD90 (5E10, BD Biosciences) markers to determine the hematopoietic-Lin-MSC fraction. Zombie Aqua^TM^ Dye (423101, BioLegend®) was used as a viability marker (1/500).

### Mice

All experimental procedures were done in accordance with the recommendations of the European Community (2010/63/UE) and French Ministry of Agriculture regulations (animal facility registration number: A9203202) for the care and use of laboratory animals. Experimental procedures on animal were approved by the French National Animal Care and Use Committee (Project A21_021, extension of Project A17_009, APAFIS#9458-2017033110277117). NOD.Cg-PrkdcscidIl2rgtm1Wjl/SzJ (NSG), originally purchased from the Jackson Laboratory (Bar Harbor, Maine, USA), were housed and bred in specific pathogen-free animal facilities (Commissariat à l’Energie Atomique et aux Energies Alternatives (CEA), Fontenay-aux-roses, France). Experiments were performed in 8 to 12-week-old NSG females. At dedicated time points, NSG mice were sub-lethally irradiated at 2Gy using a GSRD1-irradiator (137 CS source, GSM, dose rate 0.97gy/min, irradiation plateform of Institut de Radiobiology Cellulaire et Moléculaire, Fontenay-aux-Roses) 4 hours before intra-venous retro-orbital injection of human CB CD34^+^ cells under isoflurane anesthesia.

### Human Ossicles

The human ossicle (hOss) formation protocol was adapted from previous works (Reinisch et al., 2017b). Briefly, hOss were generated after subcutaneous injection of 2×10^6^ MSC. MSC were centrifuged 5 min at 300g and resuspended in 50µL human platelet lysate and the cell suspension was kept on ice. Osteogenic factor, hBMP-7 (5μg, 130-108-988, Miltenyi Biotec), was added in the mix of cells and lysate. The cell suspension was then mixed to 250µL cold matrigel (ECM625, Mercks Millipore) and the final 300µL cell mixture was injected subcutaneously in one flank of isoflurane-anesthetized mice. Every mouse received 2 injections to form 2 hOss (allowed by ethical committee rules). Bone formation was tested after injection of 4 nmol of OsteoSense 750 ex (NEV10053EX, Perkin Elmer) per mouse 24 hours before tomography analysis.

### Human hematopoietic development in NSG mice

Two to 8 weeks after hOss implantation, 1×10^5^-1×10^6^ CB CD34^+^ cells (100 µL DPBS) were injected intravenously (IV) in the sublethally irradiated mice to constitute human hematopoiesis in vivo. Three-months later, mice were sacrificed, 2 tibiae and 2 femurs (4 long bones, called mBM) and hOss were harvested from euthanized animals and hematopoietic cells were isolated by flushing the mBM with DPBS and by gently crushing hOss. The isolated cells were maintained on ice during the analysis. Immunophenotyping of human cells was performed using flow cytometry (see below). Counting was realized with a Guava Easycyte 8HT with Muse count & Viability kit. Serial transplantation was carried out after flow cytometry measurement of the percent of hCD34^+^ cells and the counting of total BM/hOss cells, by intra-venous injection of 3-5×10^5^ hCD45^+^CD34^+^ cells, recovered from primary recipient mice, in secondary sublethally irradiated animals.

### Immunostaining and coloration

Tissues were dehydrated in graded ethanol dilution baths, cleared with xylene and embedded in paraffin wax. Microtome sections (5 µm, Leica RM2245) were mounted on adhesive slides (Klinipath-KP-PRINTER ADHESIVES), deparaffinized, and stained with hematoxylin, eosin and saffran (HES) for morphology and Fast Green-Safranin O to detect proteoglycan. Slides were scanned with the Panoramic Scan 150 (3D Histech) and analyzed with the CaseCenter 2.9 viewer (3D Histech).

To visualize the vascularization of hOss, we froze directly a piece of ossicles in cryo-gel. We used CryoJane Tape-Transfert Systems of Leica Biosystems on cryotome sections of 16µm. Samples were fixed with acetone during 8 min at −20°C, dried during 3 min at RT and washed twice during 2 min and 10 min. Then a saturation was set during 30 min with normal goat serum (NGS, 15%, 005-000-121, Jackson ImmunoResearch) in PBS and blocked twice 15 min with Avidin/biotin blocking system (927301, Biolegend®). Following was a one-night incubation at 4°C with Meca32-biotin (1/250, 558773, BD Pharmingen) in PBS with 5% NGS and 0.05% Tween20. Next, primary antibodies were washed out twice during 5 min with PBS 0.1% Tween20 and 5 min with PBS. An additional incubation occurred with a Streptavidin AlexaFluor-594 (1/500, S32356, Invitrogen) diluted in PBS with 5% NGS, 0.05%Tween20 during 1h at RT following by three 5 min washes with PBS 0.1% Tween20 at RT. DAPI (1/2000) was next deposited for 5 min at RT and the samples were mounted on glass slides using Prolong Gold antifade reagent and a coverslip (P36934, Invitrogen). Samples were imaged using Spining Disk confocal microscope and analyzed by ImageJ.

Then hOSS were paraffin embedded, sectioned (4μm) and stained with MPO (Clone Poly, DAKO), human anti-CD45 antibody (clone 2B11+PD7/26, DAKO), human anti-CD34 antibody (clone QBend-10, DAKO), human anti-CD14 antibody (clone 7, BIOSB-Diagomics), human anti-Glycophorin C (clone Ret40f, DAKO), human anti-CD61 (clone 2F2, LEICA), and mouse anti-endomucin (clone V.1C7.1, Abcam). Images were acquired using a Lamina slide scanner (Perkin Elmer) and analyzed with CaseViewer software (Version 2.4.0.119028).

### Flow cytometry phenotyping of human/murine hematopoietic cells

1×10^6^ cells from mBM and from hOss were centrifuged, resuspended in 100µL DPBS and incubated with antibodies (1/100, all from BioLegend®, otherwise indicated) directed against human CD19 (HIB19), CD14 (61D3), CD15 (W6D3), CD34 (581), CD90 (5E10, BD Biosciences), CD3 (SK7), CD38 (HB-7), CD45 (HI30) and mouse TER119 (TER-119, 1/500, eBioscience), mouse CD45 (30-F11, 1/500, BD Pharmingen). Zombie Aqua™ Dye was used as a live/dead cell marker (dilution 1/500). All data were acquired on BD FACSCanto™ II and BD LSR II SORP machines with the DIVA software. Compensation controls were performed with single-stained compensation beads. After acquisition, live cells were gated and antibody-labelling analysed using FlowLogic 7 software.

### Colony Forming Units (CFU) assays

As for secondary transplantation, tested cell suspensions were characterized by flow cytometry and the percent of CD34^+^ cells carefully monitored. Based on these results, an equivalent of 1,000 human CD34^+^ cells from mBM and hOss were plated in 1 mL of methylcellulose-based medium (130-125-042, StemMACS™ HSC-CFU Assay Kit; Miltenyi Biotec) that contained human SCF, IL-3, IL-6, Erythropoietin (EPO), G-CSF, GM-CSF and 1% Penicillin/Streptomycin. Each experiment was performed in technical triplicates. The cells were incubated for 14 days at 37°C. After 14 days, human hematopoietic colonies, also called CFU, were counted and qualified by their shape and the morphology of the containing cells using an inverted microscope.

### Single cell RNA-seq

10^5^ human CD34^+^ HSPCs from a single CB source were injected intravenously into NSG mice, previously irradiated with sub-lethal doses (2Gy). Mice were either hOss-bearing or hOss-free. hOss were obtained from F/hOss (n=1) or P-N/hOss (n=2). As control condition, we used NSG mice without hOSS (mBM W/o hOss, n = 2). Three months after HSPC injection, mice were sacrificed and mononuclear cells (MNC) that had engrafted into hOSS and murine BM were pooled for each condition. In total, we had 3 samples of murine BM (mBM W/o hOss, mBM P-N/hOss and mBM F/hOss) and 2 samples of pooled ossicles (F/hOss and P-N/hOss). These 5 samples were then characterized by flow cytometry to assess i) the human engraftment (%hCD45^+^), ii) the repartition of the populations and iii) the percentage of hCD34 expressing cells (%CD45^+^/CD34^+^). To enrich samples in CD34^+^ fraction, immunomagnetic sorting was performed (130-046-703, CD34 MicroBead Kit, human, Miltenyi Biotec). After checking the purity and viability of the sorted cells, the CD34^+^ cell fraction was used to prepare scRNA-seq libraries.

### Libraries preparation

Human CD34^+^ cells were washed with PBS containing 0.04% BSA. Cell concentration and viability were determined microscopically with a Malassez counting chamber cell after staining with trypan blue. Libraries were prepared with the Chromium system of 10× Genomics, with the Chromium Single Cell 3’ v3.1 kit according to the manufacturer’s protocol (www.10xgenomics.com). The samples were processed on the same chromium chip. The number of cells targeted was 8000 per sample. Sequencing of the libraries was performed on a Miseq sequencer (Illumina) in pair-end, dual-index mode, with a target of 400 million reads per sample.

### Bioinformatic analysis

All bioinformatics analyses are based on the recommendations issued by (Luecken and Theis, 2019).

### Pre-processing of scRNA-seq data

The Galaxy interface was used to construct the gene-barcode (= Gene-cell) expression matrix. Briefly, FastQ files were aligned to the reference genome (GRCh38) via STAR (Spliced Transcripts Alignment to a Reference) (Dobin et al., 2013), with gene annotation from ENSEMBL. After quantifying the UMIs (Unique molecular identifiers) for each barcode, the unfiltered gene-barcode matrix was generated. From this unfiltered matrix, we created a filtered matrix (without empty cells and cells with a low number of UMIs (UMI < 50)). The rest of the analysis was carried out using Seurat (Version 4.1.0)316. Quality controls are based on the combined use of 3 variables: the number of UMIs per barcode, the number of genes per barcode and the fraction of reads from mitochondrial genes per barcode. We have chosen to retain for further analysis only barcodes expressing at least 200 UMI, 200 genes, with a percentage of mitochondrial genes below 20%. UMI counts were normalized with the “NormalizeData” Seurat function. A linear transformation of gene expression was then applied to prevent certain highly expressed genes from masking the expression of others (“Scaling”).

### Dimensionality reduction and Integration

For each matrix, we used the “vst” method to determine the Highly Variable Features. The selection of these genes then enabled the application of algorithms specially dedicated to dimensional reduction like PCA and UMAP. To enable comparative analysis between different experimental conditions, we then integrated data sets. Integration involves a preliminary step of identifying “anchors”, which correspond to “pairs” of biologically similar cells between the different datasets. These integrated data were then used for the graphical representations.

### Clustering and Cell annotation

Based on the UMAP generated on the integrated data, cells were grouped into clusters using the k Nearest Neighbors (KNN) algorithm. The genes most differentially expressed by each cluster were determined and used to define a list of biomarker genes. Using these lists, which we compared with publicly available data (Hay et al., 2018a), we assigned each cluster to a particular cell type. We chose not to perform cell cycle regression, so that we could study whether hOss could have an effect on the cell cycle.

### Differential gene expression

After specifying the subpopulation of interest (subset.ident) and the samples to be compared (ident.1 and identi.2), we performed a Wilcoxon test. Seurat’s “FindMarkers” function was used for these differential expression analyses. EnrichR was used for enrichment analysis.

### Barcode virus production, infection and injection

10^6^ fresh CB CD34^+^ cells/ml were pre-stimulated in BIT 9500 Serum Substitute (09500, StemCell^TM^ Technologies) supplemented with cytokines (hSCF 100ng/mL, 130-046-703; hFlt3-Ligand 100ng/mL, 130-096-479; hIL-3 60ng/mL, 130-095-069; TPO 10nM, 130-094-013, all from Miltenyi Biotec) and 1% PS during 24 hours. Sulfate protamine (5µg/mL) and 6µL barcode-containing lentivirus vector suspension were added per well on the next day. The lentiviral vector mix was produced as before (Eisele et al., 2022) by infecting HEK293T cells with the barcode plasmid, p8.9-QV and pVSVG in DMEM-GlutaMAX (Gibco) supplemented with 10% FCS (Eurobio), 1% MEM NEAA (Sigma), and 1% sodium pyruvate (Gibco) using polyethyleneimine (Polysciences). Supernatant was 0.45 μm filtered, concentrated 35 times by 1.5 hr ultracentrifugation at 31,000 × g, and frozen at –80°C. The lentiviral vector mix included a DNA stretch of 180 bp with a 20 bp ‘N’-stretch, called the barcode and a GFP fluorescent reporter. It contained around 40,000 different barcodes. CB CD34^+^ cells were spinoculated with the lentiviral particles during 1h30 at 2000rpm at room temperature and thereafter incubated during 4h30 at 37°C. The cells were then washed in αMEM media supplemented by 10% FBS, resuspended at 10^6^ cells/mL of DPBS and 10^5^ cells were injected per NSG mouse carrying or not hOss. This protocol was designed to reach a single genomic integration of the lentivector per cell, and thus the transduction was meant to be <30% efficiency that was verified by flow cytometry 72h after the end of transduction.

### Cell progeny isolation for barcoding

When clonal tracking was to be done, engrafted human hematopoietic progenitor cells (GFP^+^/hCD45^+^/CD34^+^), B cells (GFP^+^/hCD45^+^/CD19^+^) and myeloid cells (GFP^+^/CD45^+^/CD14^+^/CD15^+^) were sorted using a BD InfluxTM Cell Sorter (BD Biosciences) using antibodies described before. Sorted cells were mixed with PCR lysis buffer (301-C, Euromedex) supplemented with proteinase K solution (20mg/ml, 25530-049, Invitrogen), incubated at 55°C for 2 hours, 85°C for 30 min, 95°C for 5 min, and finally stored at 4°C.

### Lysis, barcode amplification, and sequencing

Lysed samples were then split into two replicates, and a three-step nested PCR was performed to amplify barcodes and prepare for sequencing as in (Eisele et al., 2022). In summary: the Taq DNA Polymerase Recombinant 500U (Invitrogen™) was used to perform all the PCRs. The first step amplifies barcodes (top--LIB [5′TGCTGCCGTCAACTAGA ACA-3′] and bot--LIB [5′ GATCTCGAATCAGGCGCTTA-3′]). The second step adds unique 8 bp plate indices as well as Read 1 and 2 Illumina sequences (PCR2--Read1-plate-index-partial-Top--forward 5′ ACAC TCTT TCCC TACA CGAC GCTC TTCC GATC TNNN NNN NNCTA GAAC ACTC GAGATCAG 3′ and PCR2--Read2-partial-Bot--reverse 5′ GTGA CTGG AGTT CAGA CGTG TGCT CTTCCGAT CGAT CTCG AATCAGGC GCTTA3′). In the third step, P5 and P7 flow cell attachment sequences and one of 96 sample indices of 7 bp are added (PCR3--P5-forward 5′AATGATA CGGC GACC ACCG AGAT CTAC ACTC TTTC CCTA CACG ACGC TCTT CCGATCT3′ and PCR3--P7-sample--index--reverse 5′ CAAG CAGA AGAC GGCA TACG AGAN NNNN NNGT GACT GGAG TTCAGA CGTGCTCTTCCGATC3′). (PCR program: hot start 5 min 95°C, 15 s at 95°C; 30 s at 57.2°C; 30 s at 72°C, 5 min 72°C, 4°C forever. 30 cycles [PCR1--2] or 15 cycles [PCR 3]).

Both index sequences (sample and plate) were designed based on Faircloth and Glenn, 2012 such that sequences differed by at least 2 bp. To avoid lack of diversity at the beginning of the reads, we performed the PCR2 with a mix of 4 different plate indexes for the PCR2-Read1-plate-index-partial-Top—forward primer (ACGGAATG, CTAACTCC, GATGGTCA, TGGCAGAA). Primers were ordered desalted as high-performance liquid chromatography purified. During lysis and each PCR, a mock control was added. The DNA amplification by the three PCRs was monitored by the run on a 2% agarose gel. PCR3 products for each sample and replicate were pooled, purified with the Agencourt AMPure XP system (ratio 1,2x) (Beckman Coulter) and diluted to <10nM. A High Sensitivity D1000 ScreenTape® (Agilent) was performed on the purified PCR3 pool, and sequenced on a NovaSeq system (Illumina) (SR--65bp) at the Institut Curie facility with 10% of PhiX spike-in.

### Cellular barcode sequence preprocessing

The barcodes in the NGS reads were identified by locating the variable sequence between the constant sequences to all barcodes 5’ CTAGAACACTCGAGATCAG and 3’ TGTGGTATGATGTAT using CellBarcode package (Sun et al., 2024). We only kept the barcodes that appeared in the reference list generated from sequencing the lentiviral library provided in (Eisele et al., 2022), and removed any barcodes not detected in both technical replicates. For each technical replication, we normalize the total sequencing reads to 100k reads. Then we used the mean of the normalized barcode reads from two technical replicates for further analysis. Further filtering included the removal of barcodes with less than 100 total reads across all samples. Additionally, if a barcode represented less than 1% of the reads from a specific sample compared to the total reads across all samples of that barcode, it was set to zero in the corresponding sample. To validate our filtering process, we assessed the heatmap of reads counts across samples within each batch that was generated using the heatmap package (Kolde, 2012). Finally, the read numbers were renormalized to the number of GFP positive sorted cells and any barcode with less than 2 cells were set to zero. The remaining barcodes were utilized for barcode analysis in **Figure 6B-E** and **Figure S6E-I** for which the data is renormalized to 100k reads. For the clone size analysis in **Figure 6C** and **S6F**, the data was renormalized to the total number of barcoded cells extracted during cell isolation and analyzed by cytometry to take into account the total size of the different compartments.

### Barcode analysis, plotting and statistics

We calculated the engrafted cells by dividing barcode number to estimated injected GFP positive cells (injected cell number multiplied by the fraction of GFP positive cells measured after transduction). For lineage bias analysis in **Figure S6E**, barcode clones bias towards “M” were defined as containing less than 10% of B and 10% of CD34 cells out of their total output, similarly to previous classification used in (Eisele et al., 2022). The same applies for the “B” biased clones defined as containing less than 10% of M and 10% of CD34 cells. In the analysis of compartment-specific barcodes in **Figure S6H-I**, first the count for each barcode occurring in multiple cell types within one organ were summed. Then if a barcode had zero reads in either the bone marrow or the ossicles then it was classified as under the “hOss” group or the “mBM” group respectively. The other barcodes that were present in both organs are categorized as the “both” groups. To produce the figures after filtering, R4.3.1 was used. The ternary plot was done by ggtern package (Hamilton and Ferry, 2018); boxplot and other general visualization was done with ggplot2 package (Wickham, 2011). When appropriate and as indicated in the legend, t-test was used to compare between conditions across mice, and paired t-test was used to compare conditions within mice.

### Statistical analysis

Statistical analysis was performed using GraphPad Prism version 9 software. Values are presented as the median with 95% of Cl or mean ± SEM. Statistical comparisons between conditions were determined using Kruskal-Wallis without correction or Mann-Whitney test. Differences with p<0.05 (*), p<0.01 (**), p<0.001 (***) and p<0.0001 (****) were considered statistically significant.

## Acknowledgements

We acknowledge the help of B Burroni, from the Hôpital Cochin, Paris, France (immuno-histo-chemestry labelling of hOss sections), the people from the technical platforms of Institut de Radiobiology Cellulaire et Moléculaire, IBFJ/CEA, Fontenay-aux-Roses, France, in particular V Ménard from the Irradiation Platform, D Busso and G Piton from the Molecular Bioengineering platform, J Baijer and A Schmitz from the flow cytometry platform, and J Rivière and M Vilotte from INRAE for hOss sections and histochemestry. We are grateful to F Duconge who helped with tomography and to C Antoniewski and L Bellenger from ARTbio Bioinformatics Analysis Facility from Sorbonne University. We acknowledge the fetopathology department of Antoine Béclère Hospital, Clamart, France, The clinique des Noriets, Vitry-sur-Seine, France and the Cell Therapy department from Hôpital Saint Louis, Paris, France, for providing human cord blood and BM samples. We thank the patients for agreeing to help scientific research. We thank S Teinrera Bento for help with barcode PCR in the troubleshooting phase of the project and F Cayrac from the BMBC facility for the barcode lentivirus production. We thank E Solary, N Drouin and C Jego (Gustave Roussy, Villejuif, France) and C Dussiau (Institut Cochin, Paris) who participated to the early days of the project. C Moore edited the manuscript for english language.

This work was supported by institutional grants from INSERM, CEA, Université Paris Saclay and Université Paris Cité. The study was also supported by ITMO-Cancer (3R Program, édition 2020 and MIC Program, édition 2022), CONECT-AML Pair-Pédiatrie program, Ligue Nationale Contre le Cancer (Labellisation Program) and the Fondation ARC pour la recherche contre le Cancer (ARC).

The authors declare no competing financial interests

## Authors participation

LR, CF, KG, CC, EP, VB, SD and ND performed experiments. LR, WS, CF and JC performed scRNAseq and barcoding bioinformatic analyses; KS, LK, AR, JM, AM and LF provided extremely important materials; LP, OK and FP supervised the work; LR, CF and WS made the figures; LR and FP wrote the manuscript; all authors read and approved the manuscript content.

## Notes

### Competing Interest Statement

The authors have declared no competing interest.

## References

Abarrategi, A., K. Foster, A. Hamilton, S.A. Mian, D. Passaro, J. Gribben, G. Mufti, and D. Bonnet. 2017. Versatile humanized niche model enables study of normal and malignant human hematopoiesis. J Clin Invest. 127:543–548. doi:10.1172/JCI89364.

Abarrategi, A., S.A. Mian, D. Passaro, K. Rouault-Pierre, W. Grey, and D. Bonnet. 2018. Modeling the human bone marrow niche in mice: From host bone marrow engraftment to bioengineering approaches. J Exp Med. 215:729–743. doi:10.1084/jem.20172139.

Antonelli, A., W.A. Noort, J. Jaques, B. de Boer, R. de Jong-Korlaar, A.Z. Brouwers-Vos, L. Lubbers-Aalders, J.F. van Velzen, A.C. Bloem, H. Yuan, J.D. de Bruijn, G.J. Ossenkoppele, A.C.M. Martens, E. Vellenga, R.W.J. Groen, and J.J. Schuringa. 2016. Establishing human leukemia xenograft mouse models by implanting human bone marrow-like scaffold-based niches. Blood. 128:2949–2959. doi:10.1182/blood-2016-05-719021.

Baccin, C., J. Al-Sabah, L. Velten, P.M. Helbling, F. Grünschläger, P. Hernández-Malmierca, C. Nombela-Arrieta, L.M. Steinmetz, A. Trumpp, and S. Haas. 2020. Combined single-cell and spatial transcriptomics reveal the molecular, cellular and spatial bone marrow niche organization. Nat Cell Biol. 22:38–48. doi:10.1038/s41556-019-0439-6.

Baryawno, N., D. Przybylski, M.S. Kowalczyk, Y. Kfoury, N. Severe, K. Gustafsson, K.D. Kokkaliaris, F. Mercier, M. Tabaka, M. Hofree, D. Dionne, A. Papazian, D. Lee, O. Ashenberg, A. Subramanian, E.D. Vaishnav, O. Rozenblatt-Rosen, A. Regev, and D.T. Scadden. 2019. A Cellular Taxonomy of the Bone Marrow Stroma in Homeostasis and Leukemia. Cell. 177:1915–1932.e16. doi:10.1016/j.cell.2019.04.040.

Chen, Y., R. Jacamo, Y. Shi, R. Wang, V.L. Battula, S. Konoplev, D. Strunk, N.A. Hofmann, A. Reinisch, M. Konopleva, and M. Andreeff. 2012. Human extramedullary bone marrow in mice: a novel in vivo model of genetically controlled hematopoietic microenvironment. Blood. 119:4971–4980. doi:10.1182/blood-2011-11-389957.

Cheung, A.M.S., L.V. Nguyen, A. Carles, P. Beer, P.H. Miller, D.J.H.F. Knapp, K. Dhillon, M. Hirst, and C.J. Eaves. 2013. Analysis of the clonal growth and differentiation dynamics of primitive barcoded human cord blood cells in NSG mice. Blood. 122:3129. doi:10.1182/blood-2013-06-508432.

Côme, C., A. Balhuizen, D. Bonnet, and B.T. Porse. 2020. Myelodysplastic syndrome patient-derived xenografts: from no options to many. Haematologica. 105:864–869. doi:10.3324/haematol.2019.233320.

Cordeiro Gomes, A., T. Hara, V.Y. Lim, D. Herndler-Brandstetter, E. Nevius, T. Sugiyama, S. Tani-Ichi, S. Schlenner, E. Richie, H.-R. Rodewald, R.A. Flavell, T. Nagasawa, K. Ikuta, and J.P. Pereira. 2016. Hematopoietic Stem Cell Niches Produce Lineage-Instructive Signals to Control Multipotent Progenitor Differentiation. Immunity. 45:1219–1231. doi:10.1016/j.immuni.2016.11.004.

Ding, L., and S.J. Morrison. 2013. Haematopoietic stem cells and early lymphoid progenitors occupy distinct bone marrow niches. Nature. 495:231–235. doi:10.1038/nature11885.

Dobin, A., C.A. Davis, F. Schlesinger, J. Drenkow, C. Zaleski, S. Jha, P. Batut, M. Chaisson, and T.R. Gingeras. 2013. STAR: ultrafast universal RNA-seq aligner. Bioinformatics. 29:15–21. doi:10.1093/bioinformatics/bts635.

Dupard, S.J., A. Grigoryan, S. Farhat, D.L. Coutu, and P.E. Bourgine. 2020. Development of Humanized Ossicles: Bridging the Hematopoietic Gap. Trends Mol Med. 26:552–569. doi:10.1016/j.molmed.2020.01.016.

Dykstra, B., D. Kent, M. Bowie, L. McCaffrey, M. Hamilton, K. Lyons, S.-J. Lee, R. Brinkman, and C. Eaves. 2007. Long-term propagation of distinct hematopoietic differentiation programs in vivo. Cell Stem Cell. 1:218–229. doi:10.1016/j.stem.2007.05.015.

Eisele, A.S., J. Cosgrove, A. Magniez, E. Tubeuf, S. Tenreira Bento, C. Conrad, F. Cayrac, T. Tak, A.-M. Lyne, J. Urbanus, and L. Perié. 2022. Erythropoietin directly remodels the clonal composition of murine hematopoietic multipotent progenitor cells. Elife. 11:e66922. doi:10.7554/eLife.66922.

Ergen, A.V., N.C. Boles, and M.A. Goodell. 2012. Rantes/Ccl5 influences hematopoietic stem cell subtypes and causes myeloid skewing. Blood. 119:2500–2509. doi:10.1182/blood-2011-11-391730.

Fritsch, K., S. Pigeot, X. Feng, P.E. Bourgine, T. Schroeder, I. Martin, M.G. Manz, and H. Takizawa. 2018. Engineered humanized bone organs maintain human hematopoiesis in vivo. Exp Hematol. 61:45–51.e5. doi:10.1016/j.exphem.2018.01.004.

Grigoryan, A., D. Zacharaki, A. Balhuizen, C.R. Côme, A.G. Garcia, D. Hidalgo Gil, A.-K. Frank, K. Aaltonen, A. Mañas, J. Esfandyari, P. Kjellman, E. Englund, C. Rodriguez, W. Sime, R. Massoumi, N. Kalantari, S. Prithiviraj, Y. Li, S.J. Dupard, H. Isaksson, C.D. Madsen, B.T. Porse, D. Bexell, and P.E. Bourgine. 2022. Engineering human mini-bones for the standardized modeling of healthy hematopoiesis, leukemia, and solid tumor metastasis. Sci Transl Med. 14:eabm6391. doi:10.1126/scitranslmed.abm6391.

Groen, R.W.J., W.A. Noort, R.A. Raymakers, H.-J. Prins, L. Aalders, F.M. Hofhuis, P. Moerer, J.F. van Velzen, A.C. Bloem, B. van Kessel, H. Rozemuller, E. van Binsbergen, A. Buijs, H. Yuan, J.D. de Bruijn, M. de Weers, P.W.H.I. Parren, J.J. Schuringa, H.M. Lokhorst, T. Mutis, and A.C.M. Martens. 2012. Reconstructing the human hematopoietic niche in immunodeficient mice: opportunities for studying primary multiple myeloma. Blood. 120:e9– e16. doi:10.1182/blood-2012-03-414920.

Guidi, N., M. Sacma, L. Ständker, K. Soller, G. Marka, K. Eiwen, J.M. Weiss, F. Kirchhoff, T. Weil, J.A. Cancelas, M.C. Florian, and H. Geiger. 2017. Osteopontin attenuates aging-associated phenotypes of hematopoietic stem cells. EMBO J. 36:1463. doi:10.15252/embj.201796968.

Hamilton, N.E., and M. Ferry. 2018. **ggtern**L: Ternary Diagrams Using **ggplot2**. J. Stat. Soft. 87. doi:10.18637/jss.v087.c03.

Hao, Y., S. Hao, E. Andersen-Nissen, W.M. Mauck, S. Zheng, A. Butler, M.J. Lee, A.J. Wilk, C. Darby, M. Zager, P. Hoffman, M. Stoeckius, E. Papalexi, E.P. Mimitou, J. Jain, A. Srivastava, T. Stuart, L.M. Fleming, B. Yeung, A.J. Rogers, J.M. McElrath, C.A. Blish, R. Gottardo, P. Smibert, and R. Satija. 2021. Integrated analysis of multimodal single-cell data. Cell. 184:3573–3587.e29. doi:10.1016/j.cell.2021.04.048.

Hay, S.B., K. Ferchen, K. Chetal, H.L. Grimes, and N. Salomonis. 2018a. The Human Cell Atlas bone marrow single-cell interactive web portal. Exp. Hematol. 68:51–61. doi:10.1016/j.exphem.2018.09.004.

Hay, S.B., K. Ferchen, K. Chetal, H.L. Grimes, and N. Salomonis. 2018b. The Human Cell Atlas bone marrow single-cell interactive web portal. Exp Hematol. 68:51–61. doi:10.1016/j.exphem.2018.09.004.

Henry, E., I. Souissi-Sahraoui, M. Deynoux, A. Lefèvre, V. Barroca, A. Campalans, V. Ménard, J. Calvo, F. Pflumio, and M.-L. Arcangeli. 2020. Human hematopoietic stem/progenitor cells display reactive oxygen species-dependent long-term hematopoietic defects after exposure to low doses of ionizing radiations. Haematologica. 105:2044–2055. doi:10.3324/haematol.2019.226936.

Holzapfel, B.M., D.W. Hutmacher, B. Nowlan, V. Barbier, L. Thibaudeau, C. Theodoropoulos, J.D. Hooper, D. Loessner, J.A. Clements, P.J. Russell, A.R. Pettit, I.G. Winkler, and J.-P. Levesque. 2015. Tissue engineered humanized bone supports human hematopoiesis in vivo. Biomaterials. 61:103–114. doi:10.1016/j.biomaterials.2015.04.057.

Itkin, T., S. Gur-Cohen, J.A. Spencer, A. Schajnovitz, S.K. Ramasamy, A.P. Kusumbe, G. Ledergor, Y. Jung, I. Milo, M.G. Poulos, A. Kalinkovich, A. Ludin, O. Kollet, G. Shakhar, J.M. Butler, S. Rafii, R.H. Adams, D.T. Scadden, C.P. Lin, and T. Lapi 2016. Distinct bone marrow blood vessels differentially regulate haematopoiesis. Nature. 532:323–328. doi:10.1038/nature17624.

Kolde, R. 2012. Pheatmap: pretty heatmaps. R package version. 1:726.

Lambers, F.M., F. Stuker, C. Weigt, G. Kuhn, K. Koch, F.A. Schulte, J. Ripoll, M. Rudin, and R. Müller. 2013. Longitudinal in vivo imaging of bone formation and resorption using fluorescence molecular tomography. Bone. 52:587–595. doi:10.1016/j.bone.2012.11.001.

Luecken, M.D., and F.J. Theis. 2019. Current best practices in single-cell RNA-seq analysis: a tutorial. Mol Syst Biol. 15:e8746. doi:10.15252/msb.20188746.

Maijenburg, M.W., M. Kleijer, K. Vermeul, E.P.J. Mul, F.P.J. van Alphen, C.E. van der Schoot, and C. Voermans. 2012. The composition of the mesenchymal stromal cell compartment in human bone marrow changes during development and aging. Haematologica. 97:179–183. doi:10.3324/haematol.2011.047753.

Martine, L.C., B.M. Holzapfel, J.A. McGovern, F. Wagner, V.M. Quent, P. Hesami, F.M. Wunner, C. Vaquette, E.M. De-Juan-Pardo, T.D. Brown, B. Nowlan, D.J. Wu, C.O. Hutmacher, D. Moi, T. Oussenko, E. Piccinini, P.W. Zandstra, R. Mazzieri, J.-P. Lévesque, P.D. Dalton, A.V. Taubenberger, and D.W. Hutmacher. 2017. Engineering a humanized bone organ model in mice to study bone metastases. Nat Protoc. 12:639–663. doi:10.1038/nprot.2017.002.

Miura, T., T. Kouno, M. Takano, T. Kuroda, Y. Yamamoto, S. Kusakawa, M.S. Morioka, T. Sugawara, T. Hirai, S. Yasuda, R. Sawada, S. Matsuyama, H. Kawaji, T. Kasukawa, M. Itoh, A. Matsuyama, J.W. Shin, A. Umezawa, J. Kawai, and Y. Sato. 2023. Single-Cell RNA-Seq Reveals LRRC75A-Expressing Cell Population Involved in VEGF Secretion of Multipotent Mesenchymal Stromal/Stem Cells Under Ischemia. Stem Cells Transl Med. 12:379–390. doi:10.1093/stcltm/szad029.

Passaro, D., A. Abarrategi, K. Foster, L. Ariza-McNaughton, and D. Bonnet. 2017. Bioengineering of Humanized Bone Marrow Microenvironments in Mouse and Their Visualization by Live Imaging. J Vis Exp. 55914. doi:10.3791/55914.

Pflumio, F., B. Izac, A. Katz, L.D. Shultz, W. Vainchenker, and L. Coulombel. 1996. Phenotype and function of human hematopoietic cells engrafting immune-deficient CB17-severe combined immunodeficiency mice and nonobese diabetic-severe combined immunodeficiency mice after transplantation of human cord blood mononuclear cells. Blood. 88:3731–3740.

Pinho, S., T. Marchand, E. Yang, Q. Wei, C. Nerlov, and P.S. Frenette. 2018. Lineage-Biased Hematopoietic Stem Cells Are Regulated by Distinct Niches. Dev Cell. 44:634–641.e4. doi:10.1016/j.devcel.2018.01.016.

Ramasamy, S.K., A.P. Kusumbe, T. Itkin, S. Gur-Cohen, T. Lapidot, and R.H. Adams. 2016. Regulation of Hematopoiesis and Osteogenesis by Blood Vessel-Derived Signals. Annu Rev Cell Dev Biol. 32:649–675. doi:10.1146/annurev-cellbio-111315-124936.

Reinisch, A., N. Etchart, D. Thomas, N.A. Hofmann, M. Fruehwirth, S. Sinha, C.K. Chan, K. Senarath-Yapa, E.-Y. Seo, T. Wearda, U.F. Hartwig, C. Beham-Schmid, S. Trajanoski, Q. Lin, W. Wagner, C. Dullin, F. Alves, M. Andreeff, I.L. Weissman, M.T. Longaker, K. Schallmoser, R. Majeti, and D. Strunk. 2015. Epigenetic and in vivo comparison of diverse MSC sources reveals an endochondral signature for human hematopoietic niche formation. Blood. 125:249–260. doi:10.1182/blood-2014-04-572255.

Reinisch, A., D.C. Hernandez, K. Schallmoser, and R. Majeti. 2017a. Generation and use of a humanized bone-marrow-ossicle niche for hematopoietic xenotransplantation into mice. Nat Protoc. 12:2169–2188. doi:10.1038/nprot.2017.088.

Reinisch, A., D.C. Hernandez, K. Schallmoser, and R. Majeti. 2017b. Generation and use of a humanized bone-marrow-ossicle niche for hematopoietic xenotransplantation into mice. Nat Protoc. 12:2169–2188. doi:10.1038/nprot.2017.088.

Reinisch, A., D. Thomas, M.R. Corces, X. Zhang, D. Gratzinger, W.-J. Hong, K. Schallmoser, D. Strunk, and R. Majeti. 2016. A humanized bone marrow ossicle xenotransplantation model enables improved engraftment of healthy and leukemic human hematopoietic cells. Nat Med. 22:812–821. doi:10.1038/nm.4103.

Sun, W., M. Perkins, M. Huyghe, M.M. Faraldo, S. Fre, L. Perié, and A.-M. Lyne. 2024. Extracting, filtering and simulating cellular barcodes using CellBarcode tools. Nat Comput Sci. 1–16. doi:10.1038/s43588-024-00595-7.

Thibaudeau, L., V.M. Quent, B.M. Holzapfel, A.V. Taubenberger, M. Straub, and D.W. Hutmacher. 2014. Mimicking breast cancer-induced bone metastasis in vivo: current transplantation models and advanced humanized strategies. Cancer Metastasis Rev. 33:721–735. doi:10.1007/s10555-014-9499-z.

Tikhonova, A.N., I. Dolgalev, H. Hu, K.K. Sivaraj, E. Hoxha, Á. Cuesta-Domínguez, S. Pinho, I. Akhmetzyanova, J. Gao, M. Witkowski, M. Guillamot, M.C. Gutkin, Y. Zhang, C. Marier, C. Diefenbach, S. Kousteni, A. Heguy, H. Zhong, D.R. Fooksman, J.M. Butler, A. Economides, P.S. Frenette, R.H. Adams, R. Satija, A. Tsirigos, and I. Aifantis. 2019. The bone marrow microenvironment at single-cell resolution. Nature. 569:222–228. doi:10.1038/s41586-019-1104-8.

Vaiselbuh, S.R., M. Edelman, J.M. Lipton, and J.M. Liu. 2010. Ectopic human mesenchymal stem cell-coated scaffolds in NOD/SCID mice: an in vivo model of the leukemia niche. Tissue Eng Part C Methods. 16:1523–1531. doi:10.1089/ten.tec.2010.0179.

Wickham, H. 2011. ggplot2. WIREs Computational Statistics. 3:180–185. doi:10.1002/wics.147.

