## supplemental materials for "Humanized *in vivo* bone marrow models orchestrate multi-lineage human hematopoietic cell development"

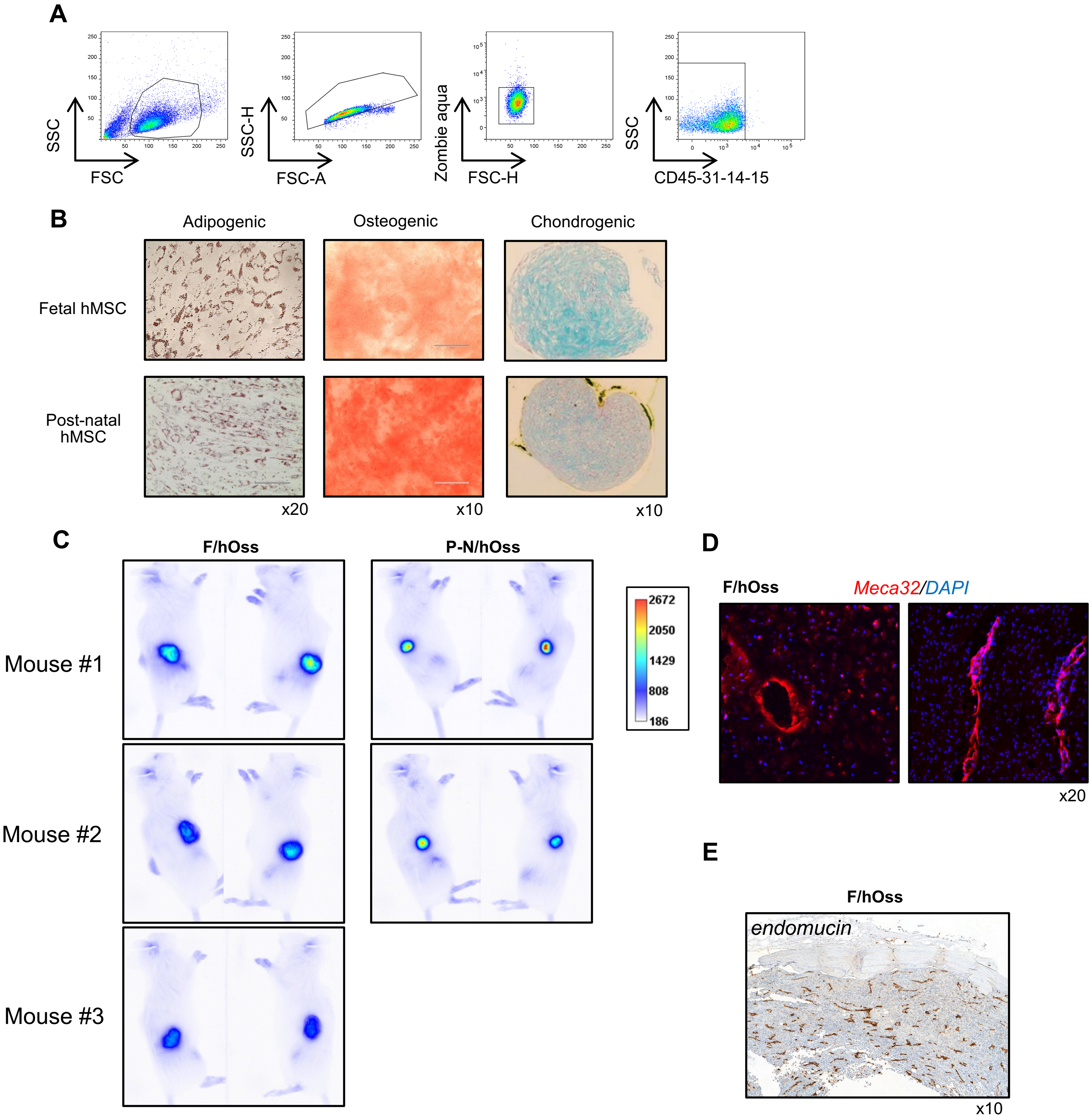


**Supplementary Figure 1:**

A. Flow cytometry gating strategy for analysis of adherent cells isolated from BM. Shown is a typical staining with anti-human CD45/CD31/CD14/CD15 (lin) antibodies and the gates used to look at CD90, CD73, CD105 and CD44 expression showing the adherent cells are lin- negative MSC (Fig1A).

B. Ex-vivo differentiation of hMSCs into adipocytes, osteoblasts et chondrocytes. N=1, 1 Fetal hBM MSC, 1 P-N hBM MSC. Shown are the results of the two hMSC tested and typical colorations used to visualize their differentiation potentials (red oil for adipocytes, red alizarin for osteocytes, and Toluidine blue for chondrocytes).

C. Measurement of bone remodeling using Osteosense and Tomography in mice with F/ (n=3) and P-N/hOss (n=2). Done at 8 weeks post-transplantation. The intensity of remodeling is pictured by a scale legend (right).

D. Immuno-fluorescence staining of F/hOss derived from the F25 (11.3 wpc) hMSC sample with anti-Meca32 antibodies and DAPI (respective red and blue staining). Shown are 2 cryostat sections of a frozen hOss (x20)

E. Immuno-histochemestry labeling of a F/hOss sample (F28) stained with anti-mouse endomucin antibody. Amplification x10.

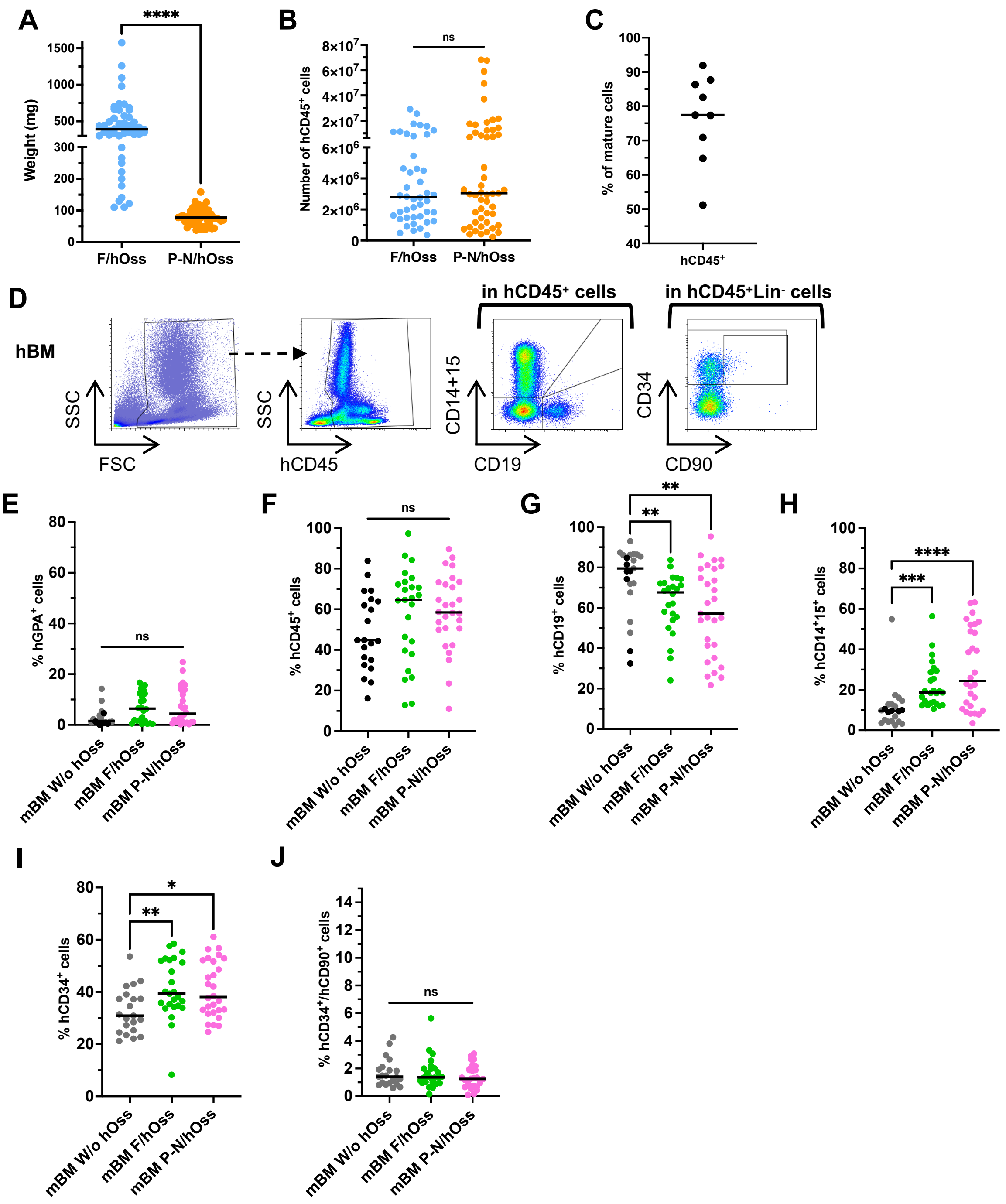


**Supplementary Figure 2.**

A. Weight of hOss obtained 12 weeks post-injection of UCB HSPC. Same mice as in Figure 2D. Median value is indicated as a line.

B. Number of hCD45⁺ cells obtained in hOss described in Figure 2E and SupFig2A. Measured 12 weeks after UCB HSPC transplant. Shown are individual values and the median (black line).

C. Percent of hCD45⁺ cells in human BM after ficoll separation. Samples used in Figure 2F-I.

D. Gating strategy used for hBM cells with data shown in Figure 2F-I.

E. Percent of human GPA⁺ erythroid in mBM from mice W/ and W/o hOss. Same gating as in Figure 2C and same mice as in Figure 2D.

F. Percent of hCD45⁺ cells detected by flow cytometry in mBM from mice w/ hOss in comparison with mBM of mice W/o hOss at 12 weeks post-transplant of UCB CD34⁺ cells. Shown are data from the same mice as in Figure 2E. Each dot identifies an individual mouse, median % is indicated by a black line. No statistical difference.

G-H. Percent of human hematopoietic cells measured in the same mice shown in Figure 2D and supFig2C. Shown are CD19⁺ B (G), CD14⁺/CD15⁺ myeloid (H)

I-J. Percent of CD34⁺ (I) and CD34⁺CD90⁺ HSC (J) gated in hCD45⁺Lin⁻ (CD19⁻CD14⁻CD15⁻) cells as shown in Figure 2C.

ns, no statistical difference; *, p<0.05; **, p<0.01; ***, p<0.001; ****, p<0.0001, (A-B) Mann-Whitney test and (E-H) Kruskal-Wallis test without correction.

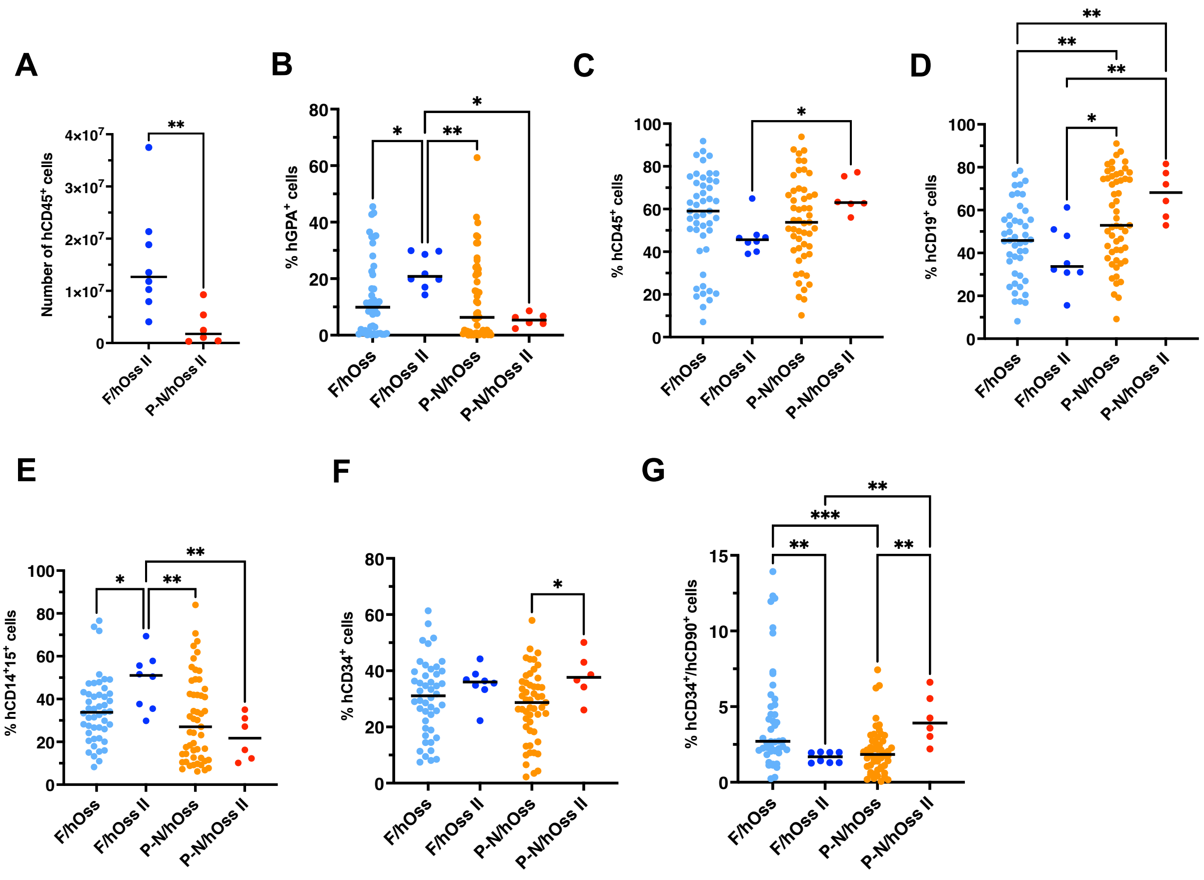


**Supplementary Figure 3.**

A. Absolute numbers of hCD45⁺ cells obtained in secondary hOss described in Figure 3G 12 weeks after CD34⁺ cell transplant. Shown are individual values of every hOss tested and the median as a black line.

B-G. Percent of human hematopoietic cells at 12 weeks post-HSPC injection but comparing primary and secondary hOss. The results are the same as the ones presented in Figure 2D-I and Figure 3F-K but put side-by-side.

ns, no statistical difference; *, p<0.05; **, p<0.01; ***, p<0.001, (A) Mann-Whitney and (B-G) Kruskal-Wallis test.

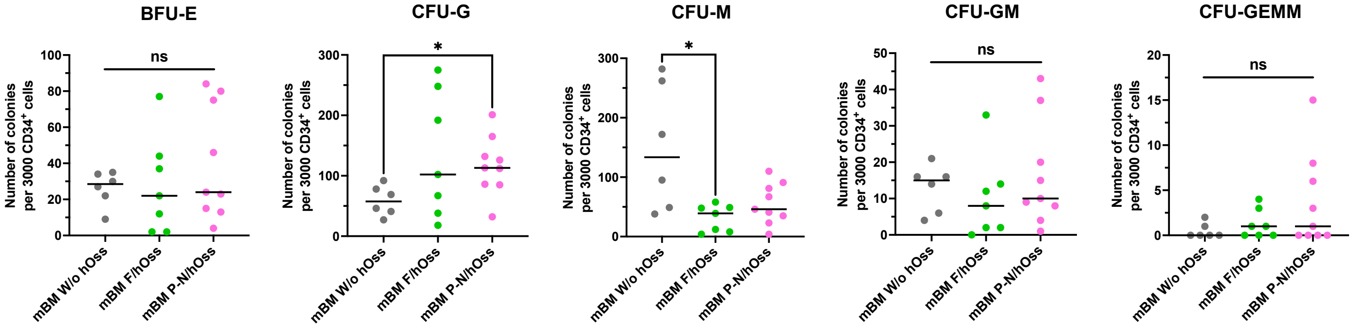


**Supplementary Figure 4.**

Numbers of every colony type generated at 2 weeks starting with 3000 hCD34⁺ cells from mBM from mice W/o (grey dots) and W/ hOss (F/hOss, green dots; P-N/hOss, pink dots). Results are from samples obtained in 3 experiments and each sample was tested in triplicates. Shown are individual means of triplicates for every tested sample. The black lines show median values. ns, no statistical difference; *, $p<0.05$, Kruskal-Wallis test without correction.

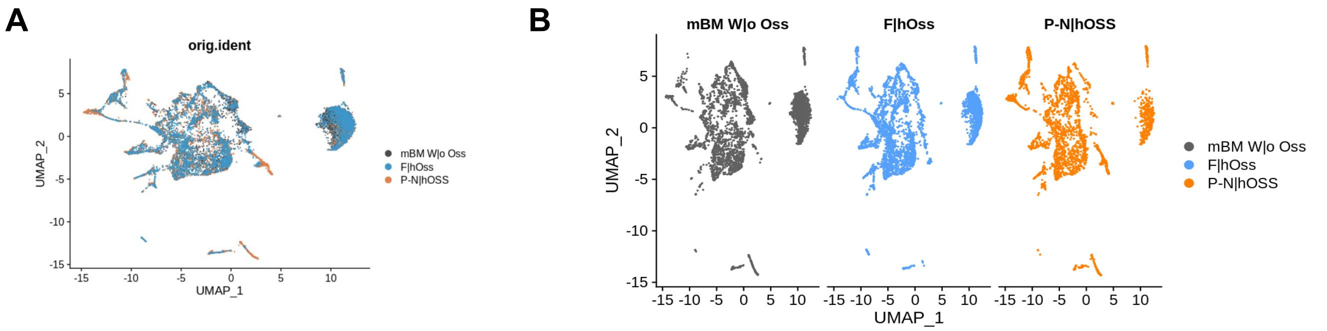


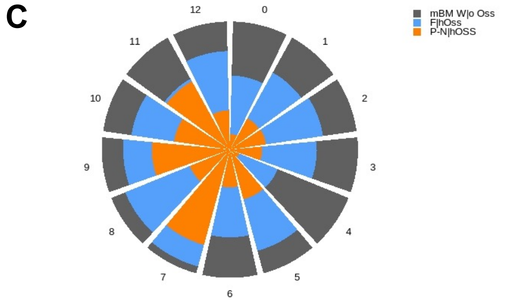


| cluster | cell population |
|---|---|
| 0 | CD34+ Pre-PC |
| 1 | CD34+ pre-B-cycling |
| 2 | CD34+ multilin/CLP |
| 3 | CD34+ HSC/MPP/LMPP |
| 4 | CD34+ pro-B |
| 5 | Immature neutrophils/monocytes |
| 6 | CD34+ MDP/Pre dendritic cells |
| 7 | CD34+ early erythroblasts |
| 8 | CD34+ Eo-B-Mast |
| 9 | Neutrophils |
| 10 | CD34+ Pre-B |
| 11 | Follicular B cells |
| 12 | Erythroblasts |

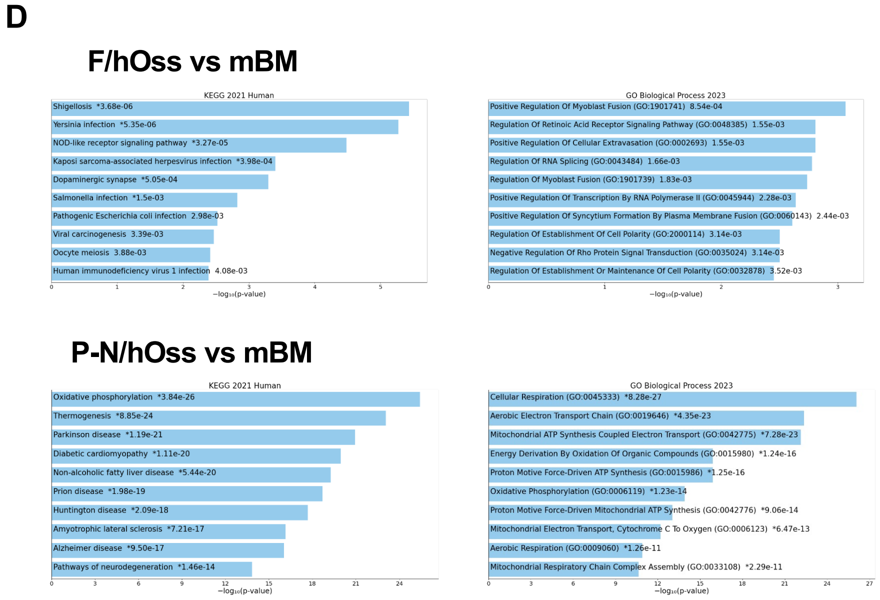


**Supplementary Figure 5.**

A. UMAP representation of the 3 integrated samples. Each point corresponds to one cell. The total number of cells is 9894.

B. UMAP of every sample analysed shown individually.

C. Repartition of cells from each sample in the different clusters, shown in proportion.

D.KEGG and GO enrichment signatures of pathways upregulated in CD34+ HSC/MPP/LMPP from P-N/hOss (B) and from F/hOss (C) compared to mBM.

**Supplemental Table 1** : Top 50 genes upregulated in P-N/hOss *vs* mBM

| gene | p_val | avg_log2FC | pct.1 | pct.2 | p_val_adj | expression |
|---|---|---|---|---|---|---|
| MIF | 3,515E-103 | 1,07421225 | 0,978 | 0,513 | 2,1319E-98 | UP |
| KRTCAP2 | 2,9928E-57 | 0,28732934 | 0,737 | 0,149 | 1,8151E-52 | UP |
| COX7A2 | 4,7536E-54 | 0,48281579 | 0,988 | 0,769 | 2,883E-49 | UP |
| MTATP6P1 | 8,8995E-53 | 0,47918439 | 0,968 | 0,638 | 5,3974E-48 | UP |
| COX6B1 | 2,6195E-51 | 0,47436822 | 0,995 | 0,869 | 1,5887E-46 | UP |
| RPL36AL | 2,183E-50 | 0,47606131 | 0,983 | 0,803 | 1,324E-45 | UP |
| IFITM1 | 1,2601E-48 | 0,19265698 | 0,491 | 0,026 | 7,6425E-44 | UP |
| COX5B | 1,0853E-47 | 0,44360304 | 0,978 | 0,715 | 6,5823E-43 | UP |
| COX6C | 5,2347E-47 | 0,44669187 | 0,988 | 0,856 | 3,1748E-42 | UP |
| RPL41P2 | 6,4959E-43 | 0,26196241 | 0,759 | 0,259 | 3,9397E-38 | UP |
| NDUFA4 | 1,9968E-41 | 0,38011747 | 0,993 | 0,8 | 1,2111E-36 | UP |
| COMMD6 | 9,9337E-38 | 0,36860728 | 0,968 | 0,703 | 6,0247E-33 | UP |
| OST4 | 1,2323E-37 | 0,36302437 | 0,99 | 0,877 | 7,4735E-33 | UP |
| HLA-C | 1,3921E-37 | 0,24152107 | 0,74 | 0,256 | 8,4432E-33 | UP |
| S100A4 | 1,8053E-36 | 0,45150509 | 0,975 | 0,813 | 1,0949E-31 | UP |
| NDUFS5 | 2,6399E-36 | 0,3673322 | 0,929 | 0,628 | 1,6011E-31 | UP |
| TXN | 2,6106E-35 | 0,37366606 | 0,958 | 0,751 | 1,5833E-30 | UP |
| RP11-770J1.8 | 3,9096E-34 | 0,15885322 | 0,592 | 0,136 | 2,3711E-29 | UP |
| RBX1 | 7,1769E-34 | 0,30673319 | 0,867 | 0,492 | 4,3527E-29 | UP |
| NDUFB2 | 2,0412E-33 | 0,35930915 | 0,934 | 0,674 | 1,238E-28 | UP |
| TOMM5 | 2,5294E-33 | 0,22194497 | 0,683 | 0,249 | 1,5341E-28 | UP |
| NDUFA13 | 6,7968E-33 | 0,29253144 | 0,904 | 0,549 | 4,1222E-28 | UP |
| MZT2B | 5,8332E-32 | 0,32204331 | 0,968 | 0,792 | 3,5378E-27 | UP |
| GMFG | 6,507E-32 | 0,32135343 | 0,973 | 0,749 | 3,9464E-27 | UP |
| COX8A | 7,8739E-31 | 0,32072738 | 0,916 | 0,61 | 4,7754E-26 | UP |
| SEC61B | 1,1433E-30 | 0,3007033 | 0,897 | 0,587 | 6,9338E-26 | UP |
| COX7B | 1,0812E-29 | 0,37806206 | 0,966 | 0,697 | 6,5573E-25 | UP |
| UXT | 1,0892E-29 | 0,28884164 | 0,956 | 0,626 | 6,6058E-25 | UP |
| SEM1 | 1,3189E-29 | 0,31652314 | 0,966 | 0,805 | 7,9993E-25 | UP |
| RPL22L1 | 1,7293E-29 | 0,43437914 | 0,966 | 0,728 | 1,0488E-24 | UP |
| NDUFS6 | 2,0894E-29 | 0,29747701 | 0,885 | 0,559 | 1,2672E-24 | UP |
| NDUFB7 | 2,6188E-29 | 0,2381354 | 0,83 | 0,441 | 1,5883E-24 | UP |
| UQCRH | 7,8292E-29 | 0,321767 | 0,988 | 0,841 | 4,7483E-24 | UP |
| ELOB | 8,5733E-29 | 0,31351622 | 0,978 | 0,833 | 5,1996E-24 | UP |
| SNRPE | 9,6757E-29 | 0,34961376 | 0,985 | 0,815 | 5,8682E-24 | UP |
| SNRPF | 1,0515E-28 | 0,37781916 | 0,983 | 0,867 | 6,3773E-24 | UP |
| NOP10 | 1,3866E-28 | 0,25068672 | 0,86 | 0,479 | 8,4097E-24 | UP |
| UBL5 | 5,0589E-27 | 0,34579641 | 0,998 | 0,897 | 3,0682E-22 | UP |
| NDUFB4 | 1,5761E-26 | 0,28299808 | 0,924 | 0,685 | 9,559E-22 | UP |
| LSM7 | 2,7394E-26 | 0,2720479 | 0,892 | 0,595 | 1,6614E-21 | UP |
| NDUFB3 | 5,2734E-26 | 0,2345175 | 0,816 | 0,467 | 3,1982E-21 | UP |
| LRRC75A | 8,0446E-26 | 0,40973611 | 0,713 | 0,382 | 4,879E-21 | UP |
| SNRPG | 1,2144E-25 | 0,33399638 | 0,948 | 0,759 | 7,3654E-21 | UP |
| NDUFB11 | 1,6152E-25 | 0,25625522 | 0,946 | 0,677 | 9,7962E-21 | UP |
| ATP5MF | 5,9601E-25 | 0,33972326 | 0,963 | 0,828 | 3,6147E-20 | UP |
| ATP5IF1 | 1,1525E-24 | 0,27611226 | 0,943 | 0,692 | 6,9901E-20 | UP |
| NME2 | 1,2109E-24 | 0,09748642 | 0,403 | 0,079 | 7,3437E-20 | UP |
| SMIM26 | 1,5197E-24 | 0,19084849 | 0,794 | 0,408 | 9,217E-20 | UP |
| C19orf53 | 2,6212E-24 | 0,204162 | 0,806 | 0,436 | 1,5898E-19 | UP |

**Supplemental Table 2** : Top 50 genes upregulated in F/hOss *vs* mBM

| gene | p_val | avg_log2FC | pct.1 | pct.2 | p_val_adj | expression |
|---|---|---|---|---|---|---|
| LRRC75A | 2,2638E-41 | 1,24422807 | 0,628 | 0,382 | 1,373E-36 | UP |
| SERF2 | 2,9269E-19 | 0,46252412 | 0,854 | 0,967 | 1,7751E-14 | UP |
| HINT1 | 2,2227E-18 | 0,46881288 | 0,827 | 0,938 | 1,348E-13 | UP |
| FTL | 5,119E-18 | 0,4603151 | 0,846 | 0,954 | 3,1046E-13 | UP |
| RPS10 | 1,4507E-17 | 0,40338694 | 0,869 | 0,974 | 8,7981E-13 | UP |
| STMN1 | 2,1382E-17 | 0,63685611 | 0,742 | 0,867 | 1,2968E-12 | UP |
| RSBN1L | 2,3254E-17 | 0,00042619 | 0,177 | 0,503 | 1,4103E-12 | UP |
| ODF2L | 4,93E-15 | 0,00760561 | 0,215 | 0,541 | 2,99E-10 | UP |
| GNG11 | 5,0946E-15 | 0,00822446 | 0,24 | 0,585 | 3,0898E-10 | UP |
| ITSN2 | 8,3814E-15 | 0,00728123 | 0,241 | 0,605 | 5,0832E-10 | UP |
| TBCA | 8,9142E-15 | 0,44537082 | 0,824 | 0,946 | 5,4064E-10 | UP |
| YWHAE | 2,0382E-14 | 0,4219753 | 0,81 | 0,941 | 1,2361E-09 | UP |
| CD47 | 3,5256E-14 | 0,00603488 | 0,272 | 0,644 | 2,1382E-09 | UP |
| CBX3 | 5,2695E-14 | 0,4843884 | 0,837 | 0,956 | 3,1959E-09 | UP |
| ZNF37BP | 5,6976E-14 | 0,01148549 | 0,168 | 0,444 | 3,4555E-09 | UP |
| HELZ | 8,3248E-14 | 0,02201156 | 0,191 | 0,49 | 5,0489E-09 | UP |
| SUPT6H | 1,0085E-13 | 0,00876898 | 0,143 | 0,39 | 6,1164E-09 | UP |
| ATP5MJ | 1,1739E-13 | 0,41996936 | 0,737 | 0,887 | 7,1197E-09 | UP |
| NFKB1 | 2,3084E-13 | 0,01055052 | 0,154 | 0,415 | 1,4E-08 | UP |
| UBAP2 | 2,9597E-13 | 0,01204393 | 0,191 | 0,477 | 1,795E-08 | UP |
| ITPR1 | 3,1671E-13 | 0,00968934 | 0,232 | 0,556 | 1,9208E-08 | UP |
| PSME4 | 3,1804E-13 | 0,00228715 | 0,21 | 0,523 | 1,9289E-08 | UP |
| EPS15 | 3,5834E-13 | 0,01558358 | 0,126 | 0,356 | 2,1733E-08 | UP |
| NORAD | 3,6702E-13 | 0,00316342 | 0,204 | 0,505 | 2,2259E-08 | UP |
| SFT2D2 | 4,2652E-13 | 0,02004697 | 0,151 | 0,4 | 2,5868E-08 | UP |
| HSD17B4 | 4,4402E-13 | 0,01021825 | 0,17 | 0,436 | 2,693E-08 | UP |
| PPP1CC | 4,5943E-13 | 0,00371604 | 0,199 | 0,49 | 2,7864E-08 | UP |
| ATG14 | 4,6133E-13 | 0,01160538 | 0,109 | 0,321 | 2,7979E-08 | UP |
| ATP5MC2 | 5,13E-13 | 0,37145791 | 0,835 | 0,959 | 3,1113E-08 | UP |
| TIMM10 | 5,5111E-13 | 0,00150874 | 0,173 | 0,441 | 3,3424E-08 | UP |
| ARF6 | 6,1154E-13 | 0,00445371 | 0,201 | 0,492 | 3,7089E-08 | UP |
| CTBP2 | 7,69E-13 | 0,01008008 | 0,177 | 0,454 | 4,6639E-08 | UP |
| EXOC1 | 8,4549E-13 | 0,02044508 | 0,202 | 0,497 | 5,1278E-08 | UP |
| DPM3 | 9,1129E-13 | 0,00870368 | 0,212 | 0,513 | 5,5269E-08 | UP |
| SEC31A | 9,9491E-13 | 0,01380696 | 0,207 | 0,505 | 6,034E-08 | UP |
| RP11-566K19.6 | 1,1084E-12 | 0,01292943 | 0,118 | 0,338 | 6,7225E-08 | UP |
| UBALD2 | 1,1418E-12 | 0,00128182 | 0,084 | 0,267 | 6,9251E-08 | UP |
| PKN2 | 1,1833E-12 | 0,01466189 | 0,188 | 0,469 | 7,1764E-08 | UP |
| ITPRIPL2 | 1,1913E-12 | 0,0061641 | 0,082 | 0,264 | 7,2252E-08 | UP |
| MAP7D1 | 1,2258E-12 | 0,00265719 | 0,09 | 0,282 | 7,4341E-08 | UP |
| RNU4-2 | 1,2281E-12 | 0,00397692 | 0,107 | 0,315 | 7,4482E-08 | UP |
| SAMD9 | 1,2799E-12 | 0,00054072 | 0,086 | 0,272 | 7,7627E-08 | UP |
| RRP1B | 1,2823E-12 | 0,00311604 | 0,19 | 0,469 | 7,7769E-08 | UP |
| ZNF326 | 1,5446E-12 | 0,00799869 | 0,235 | 0,551 | 9,3676E-08 | UP |
| ZEB1 | 1,5512E-12 | 0,00492284 | 0,121 | 0,336 | 9,4077E-08 | UP |
| KDM1B | 1,5953E-12 | 0,00743467 | 0,09 | 0,282 | 9,6752E-08 | UP |
| ELMO1 | 1,7656E-12 | 0,006844 | 0,241 | 0,559 | 1,0708E-07 | UP |
| MAPK14 | 1,7964E-12 | 0,00739431 | 0,138 | 0,369 | 1,0895E-07 | UP |
| SMIM24 | 1,8175E-12 | 0,46624344 | 0,776 | 0,91 | 1,1023E-07 | UP |

**Supplemental Table 3**: List of upregulated genes common to F/hOss vs mBM and P-N/hOss vs mBM

| **47 Upregulated Genes** | | | | | |
|---|---|---|---|---|---|
| C4orf3 | COX14 | TMEM18 | OXLD1 | LRRC75A | GNG5 |
| COX17 | SNX10 | DPM3 | SLC25A13 | LAMTOR5 | MICOS13 |
| BLOC1S1 | PTTG1IP | ADSS2 | PTPN1 | ANAPC11 | NDUFB7 |
| SMIM26 | RBMX2 | TCEA1 | TRIM52-AS1 | CHCHD10 | EIF2AK4 |
| RNF181 | UBE2V1 | SNRPB2 | SMARCE1 | TMEM256 | H1-2 |
| C9orf16 | PBDC1 | NDUFA2 | COX7A2 | TIMM10 | VPS4B |
| LSM7 | PPP1CA | NDUFB3 | PKN2 | LAMTOR4 | PAXBP1 |
| TDG | LPIN2 | MRPS21 | STAMBP | PTRHD1 | |

In green: genes found in the 50 most-upregulated genes in the two differential list (see Figure 5I)
In red: common gene found among the most upregulated 50 genes (Figure 5I)

**Supplemental Table 4**: Gene sets tested for enrichment and GSEA

| Gene set |
|---|
| BIOCARTA_CELLCYCLE_PATHWAY.v2022.1.Hs |
| FISCHER_G1_S_CELL_CYCLE.v2022.1.Hs |
| FISCHER_G2_M_CELL_CYCLE.v2022.1.Hs |
| GOBP_CELL_CYCLE.v2022.1.Hs |
| GOBP_CELL_CYCLE_PROCESS.v2022.1.Hs |
| GOBP_G0_TO_G1_TRANSITION.v2022.1.Hs |
| GOBP_HEMATOPOIETIC_STEM_CELL_DIFFERENTIATION.v7.5.1(1) |
| GOBP_HEMATOPOIETIC_STEM_CELL_HOMEOSTASIS.v7.5.1(1) |
| GOBP_HEMATOPOIETIC_STEM_CELL_MIGRATION.v7.5.1(1) |
| GOBP_HEMATOPOIETIC_STEM_CELL_MIGRATION_TO_BONE_MARROW.v7.5.1 |
| GOBP_HEMATOPOIETIC_STEM_CELL_PROLIFERATION.v7.5.1 |
| GOBP_POSITIVE_REGULATION_OF_HEMATOPOIETIC_STEM_CELL_PROLIFERATION.v7.5.1 |
| GOBP_REGULATION_OF_HEMATOPOIETIC_STEM_CELL_DIFFERENTIATION.v7.5.1 |
| GOBP_REGULATION_OF_HEMATOPOIETIC_STEM_CELL_PROLIFERATION.v7.5.1 |
| IVANOVA_HEMATOPOIESIS_STEM_CELL.v7.5.1 |
| IVANOVA_HEMATOPOIESIS_STEM_CELL_AND_PROGENITOR.v7.5.1 |
| IVANOVA_HEMATOPOIESIS_STEM_CELL_LONG_TERM.v7.5.1 |
| IVANOVA_HEMATOPOIESIS_STEM_CELL_SHORT_TERM.v7.5.1 |
| JAATINEN_HEMATOPOIETIC_STEM_CELL_DN.v7.5.1 |
| JAATINEN_HEMATOPOIETIC_STEM_CELL_UP.v7.5.1 |
| WP_HEMATOPOIETIC_STEM_CELL_DIFFERENTIATION.v7.5.1 |

### Supplementary Table 5

| Mouse ID | Analysed cells are from | Transduction efficiency (%) | Nb bc | Engrafted Stem Cell Frequency |
|---|---|---|---|---|
| **C506-S4** | W/o hOss | 7.8 | 6 | 0.001 |
| **C506-S5** | W/o hOss | 7.8 | 17 | 0.002 |
| **C506-S2** | F/hOss | 7.8 | 9 | 0.001 |
| **C506-S3** | F/hOss | 7.8 | 29 | 0.004 |
| **C342-S3** | F/hOss | 13.51 | 28 | 0.002 |
| **C507-S1** | W/o hOss | 7.8 | 8 | 0.001 |
| **C507-S3** | P-N/hOss | 7.8 | 27 | 0.003 |
| **C507-S4** | P-N/hOss | 7.8 | 13 | 0.002 |
| **C463-S4** | W/o hOss | 16.8 | 59 | 0.004 |
| **C463-S5** | W/o hOss | 16.8 | 112 | 0.007 |
| **C463-S2** | P-N/hOss | 16.8 | 186 | 0.011 |
| **C463-S3** | P-N/hOss | 16.8 | 151 | 0.009 |

### Supplemental figure 6

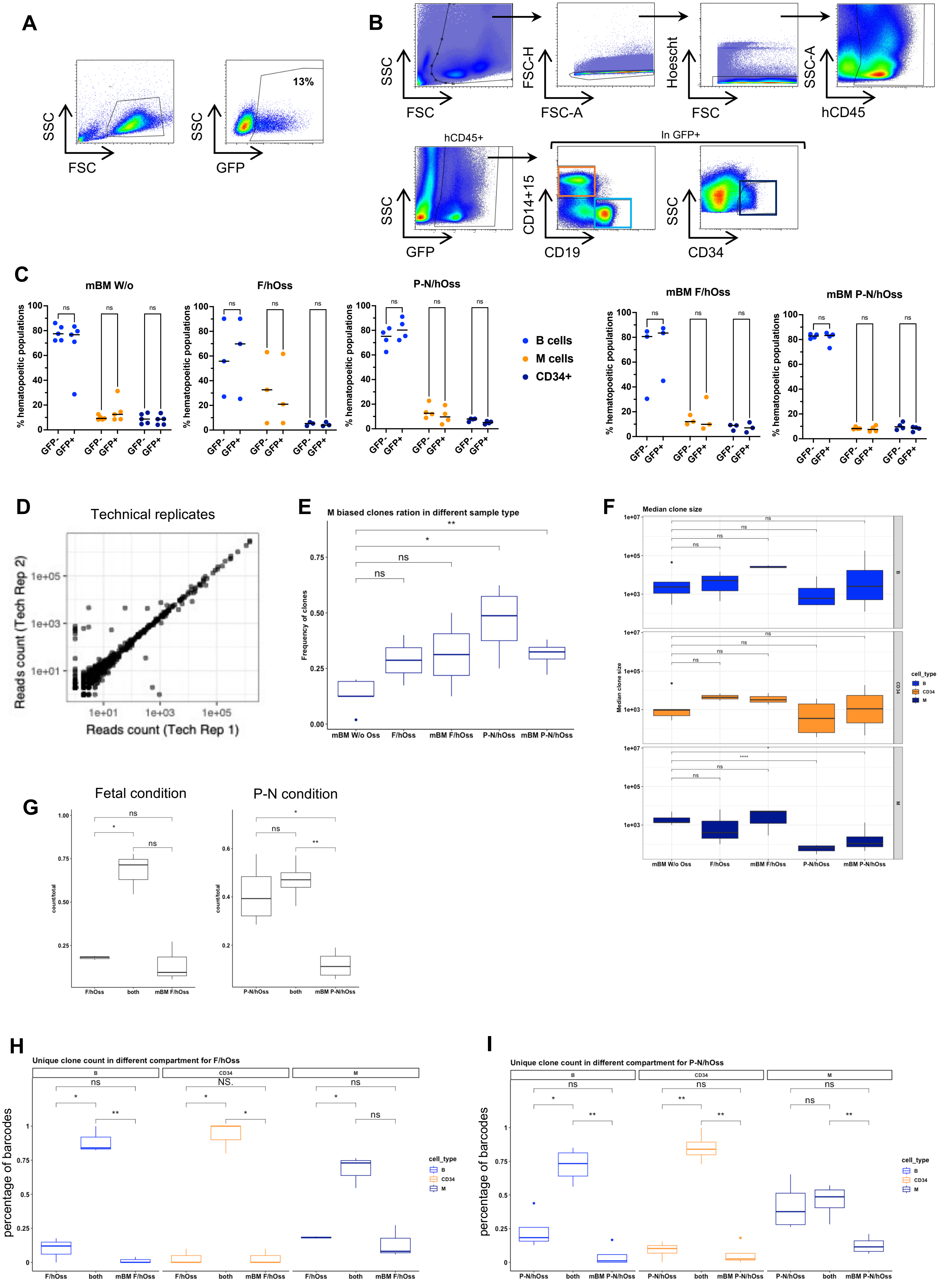

**Supplementary Figure 6.**

A. Example FACS plot (% of GFP) after transduction

B. FACS gating to define each cell type and sorted cells.

C. Percentage of $GFP^+$ and $GFP^-$ in each cell type.

D. Representative example of the barcode read counts between the technical duplicates after filtering barcodes using the reference list only. The data was renormalized to 100k reads per sample. Each point represents a barcode.

E. Percentage of M biased barcode clone between mice bearing no ossicle, F/hOss or P-N/hOss. M biased clone is defined as a barcoded clone having less than 1% of B and less than 1% of $CD34^+$ of the total number of cells produced. Clone numbers were compared between groups using t-test.

F. Median clone size among barcode clones in each BM sites of each mice per each condition. Clone sizes were compared between groups using t-test.

G. Proportion of unique clones present in both mBM and hOss ("both") or in one organ only bone marrows (mBM) or ossicles (hOss). The proportions were calculated as the number of barcodes present in one of the 3 categories (both, mBM or Oss) divided by the total number of barcodes. Clone numbers were compared using paired t-test.

H-I. The proportion of unique clones present in both mBM and hOss ("both") or in one organ only bone marrow (mBM) or ossicle (hOss)) for each lineage (B, CD34, M) analyzed in mice with FtOss (H) or PnOss(I). The proportions were calculated as the number of barcodes present in one of the 3 categories (both, mBM or Oss) divided by the total number of barcodes. Clone numbers were compared using paired t-test.

ns, no statistical difference; *, $p<0.05$; **, $p<0.01$; ***. Mann and Withney Test (C) and t-test (E-I)
